# Detection of new pioneer transcription factors as cell-type specific nucleosome binders

**DOI:** 10.1101/2023.05.10.540098

**Authors:** Yunhui Peng, Wei Song, Vladimir B. Teif, Ivan Ovcharenko, David Landsman, Anna R. Panchenko

**Affiliations:** Institute of Biophysics and Department of Physics, Central China Normal University, Wuhan 430079, China; National Library of Medicine, National Institutes of Health, Bethesda, MD, USA; School of Life Sciences, University of Essex, Wivenhoe Park, Colchester, CO4 3SQ, UK; Department of Pathology and Molecular Medicine, Queen’s University, ON, Canada; Department of Biology and Molecular Sciences, Queen’s University, ON, Canada; School of Computing, Queen’s University, ON, Canada; Ontario Institute of Cancer Research, Toronto, ON, Canada

## Abstract

Wrapping of DNA into nucleosomes restricts accessibility to the DNA and may affect the recognition of binding motifs by transcription factors. A certain class of transcription factors, the pioneer transcription factors, can specifically recognize their DNA binding sites on nucleosomes, may initiate local chromatin opening and facilitate the binding of co-factors in a cell-type-specific manner. For the majority of human pioneer transcription factors, the locations of their binding sites, mechanisms of binding and regulation remain unknown. We have developed a computational method to predict the cell-type-specific ability of transcription factors to bind nucleosomes by integrating ChIP-seq, MNase-seq and DNase-seq data with details of nucleosome structure. We have demonstrated the ability of our approach in discriminating pioneer from canonical transcription factors and predicted new potential pioneer transcription factors in H1, K562, HepG2 and HeLa cell lines. Lastly, we systemically analyzed the interaction modes between various pioneer transcription factors and detected several clusters of distinctive binding sites on nucleosomal DNA.

## Introduction

In eukaryotic cell, DNA is packaged in the form of chromatin yet, it should be dynamically accessed during transcription and replication processes at high spatiotemporal precision^1^. Open chromatin is thought to comprise actively transcribed genes, while compact chromatin contains repressed genes. However, many recent observations point to a limited association between DNA accessibility, chromatin compaction and gene transcription at the global genomic scale. Indeed, rapid transcription activation may occur by relatively small changes of DNA solvent exposure, and localized nucleosomal array structures can be dynamically exposed without the large-scale chromatin rearrangements^2–5^. Nucleosomes represent the basic subunits of chromatin structure and function. They comprise a histone octamer of four types of core histones of two copies each, and ∼147 base pairs of DNA wrapped around them^6^. Wrapping of DNA into nucleosomes inherently restricts DNA accessibility and the recognition of binding motifs by transcription factors. Intrinsically disordered histone tails flank histone core domains and may also modulate DNA accessibility by forming transient interactions with the nucleosomal and linker DNA^7^. The control of the DNA accessibility at nucleosomal and subnucleosomal scales is of major importance in understanding of how certain transcription factors can target compact chromatin to induce transcription activation or repression^8–10^.

The differentiation of cells into different lineages occurs through chromatin reprogramming, involving the cooperative behavior of various transcription factors^11,12^. Although nucleosomes generally hinder the binding of transcription factors (TFs), a certain class, so-called pioneer transcription factors (PTF), can specifically recognize their binding sites on nucleosomal DNA, in some cases initiating local chromatin opening and facilitating subsequent binding of other co-factors in a cell-type specific manner^2,12^. Several studies have revealed the critical roles of pioneer transcription factors in mediating the cell-type specific gene expression and establishment of cell lineage reprogramming^11–13^.

Significant efforts have been made to characterize the interaction landscape between various TFs and nucleosomes^14–18^. Using the NCAP–SELEX approach, a recent study has characterized the interaction modes between nucleosomes and 220 transcription factors^14^. Another high-throughput protein microarray study of 593 human TFs systematically identified the structural features of TFs binding with nucleosomes ^19^. It has been revealed that the vast majority of TFs preferably bind naked DNA instead of nucleosomal DNA at physiological concentrations, whereas certain transcription factors specifically target nucleosomes at different locations and orientations^14–16^. Moreover, several structures of pioneer transacription factors in complex with nucleosomes have recently been resolved^20–23^. Despite significant advances in recent experimental approaches, the interaction modes of most transcription factors with nucleosomes remain obscure and their cell-type-specific pioneer activities are largely unknown. The development of new computational approaches can improve our understanding of binding properties of various TFs with nucleosomes, thereby helping to identify novel pioneer transcription factors.

Through the advances in high-throughput sequencing techniques, a large volume of data has been generated (*e.g.* ChIP–seq, ATAC-seq, Dnase-seq and MNase-seq data^24^), which allows us to gain insights into the chromatin structure and details of epigenetic binding events. The rapid growth of such data sets has further stimulated the development of computational methods to characterize cell-type specific transcription factors ^25–28^. Several machine learning models have been proposed to predict transcription factor binding sites and identify sequence context features critical for TF binding^25,27–29^. In addition, gene regulatory network-based approaches have helped identify the key transcription factors in cell fate determination^30,31^. Despite the success of computational methods in genome-wide prediction of binding sites of canonical transcription factors, the locations of pioneer transcription factors’ binding sites, mechanisms of binding and regulation have not been systematically explored.

Integrating data on nucleosome positioning and DNA accessibility (MNase-seq, ATAC-seq and DNase-seq) with the data on DNA binding events, available through ChIP-seq and other methods, can reveal the interplay between transcription factor binding and nucleosome positioning, providing insights into the mechanisms of pioneer transcription factor interactions with nucleosomes and their ability to modulate chromatin accessibility^32–35^.We have developed a computational method to study the ability of transcription factors to bind nucleosomes by using ChIP-seq, MNaseq-seq and DNase-seq data sets from five different cell lines and. Our results point to the capability of our method to discriminate between pioneer and canonical transcription factors using experimental benchmarks. Additionally, we have predicted several transcription factors as potentially new cell-type specific pioneer transcription factors in H1, K562, HepG2 and HeLa cell lines and performed multiple validations. Lastly, we systemically analyzed the interaction modes between various pioneer transcription factors and nucleosomes and detected six clusters of distinctive binding sites on nucleosomal DNA.

## Methods

### Genome-wide mapping of nucleosome dyads and footprint regions

The overall workflow of our computational framework is shown in Figure 1. High-coverage micrococcal nuclease sequencing data (MNase-seq) of five human cell lines (H1, HepG2, MCF-7, K562, and HeLa) from paired-end sequencing was used for nucleosome mapping (Supplementary Table 1). The raw MNase-seq data was downloaded from the NCBI Sequence Read Archive (SRA) and converted into the fastq format using SRA Toolkit^36^. Then, the downloaded fastq files were processed using the mnaseseq pipeline from nf-core (https://github.com/nf-core/mnaseseq)^37^, a recently developed bioinformatics pipeline for MNase seq data analysis. Adapter trimming of sequencing reads was performed with Trim Galore. Then, adapter-trimmed reads were mapped to the reference genome using Burrows-Wheeler Aligner (BWA)^38^. Human genome GRCh37 was used as a reference genome for reads mapping and the maximum number of mismatches in alignment was set to 4. The minimum and maximum insert sizes for filtering of mono-nucleosome paired-end reads were set to 120 and 180 base pairs. Duplicate reads were marked using Picard MarkDuplicates command (http://broadinstitute.github.io/picard/) and discarded from the analysis to avoid PCR duplication artifacts (less than 10% of reads were duplicated). Read libraries of replicates from the same experiment condition were merged into the analysis. Next, the BAM sequence alignment files were converted into BED format using bedtools v2.30.0^39^. The aligned reads with fragment sizes from 146 to 148 base pairs were selected for mapping of the representative dyad positions (center of the nucleosomal DNA).

**Figure 1.**
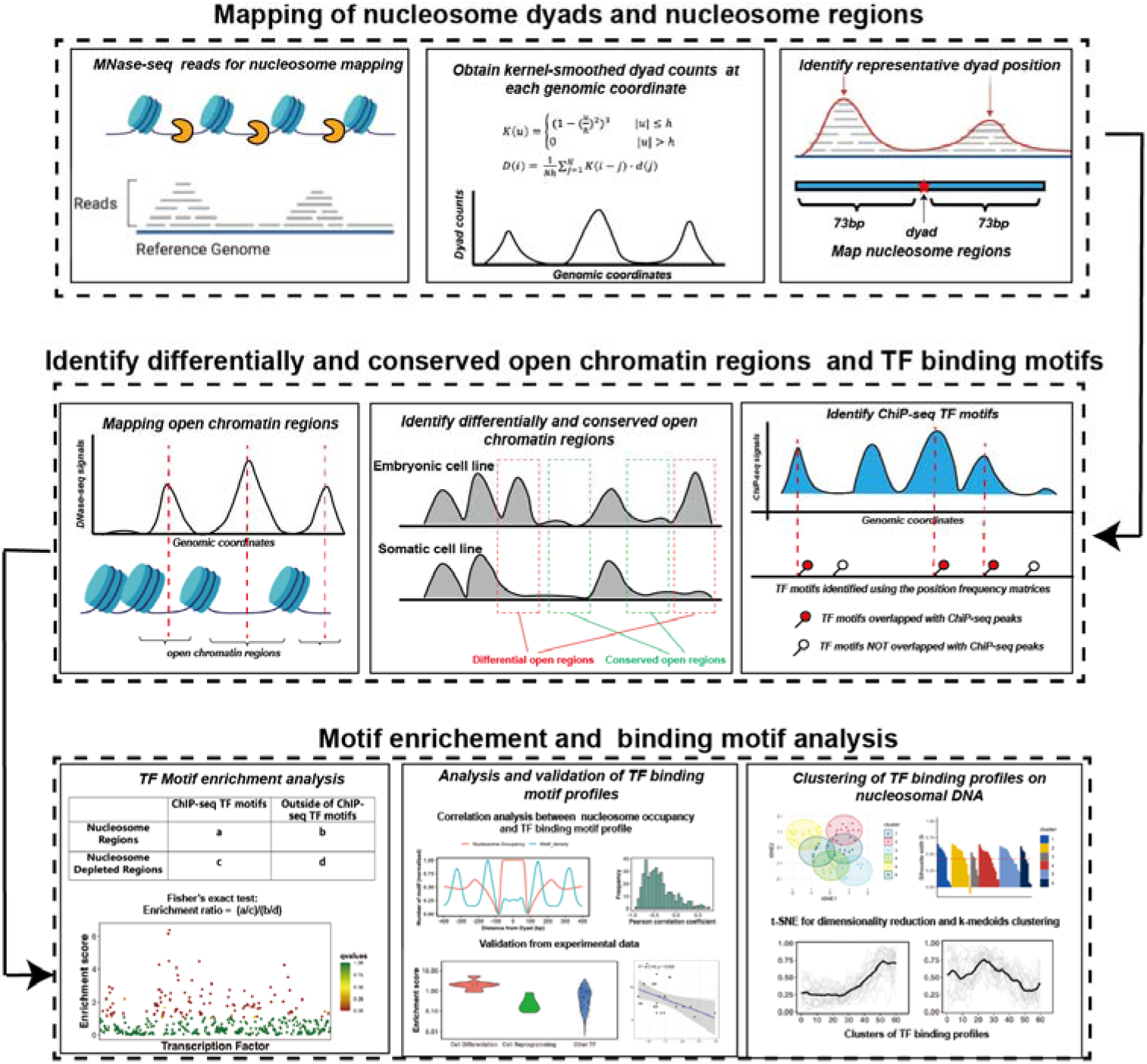
The computational framework to analyze the ability of transcription factors to bind to nucleosomes by integrating ChIP-seq, MNaseq-seq and DNase-seq data for motif enrichment and binding motif analysis.

To determine the representative dyad position of nucleosomes, we implemented and modified a previously developed nucleosome mapping protocol^40^ and dyad positions were determined as midpoints of mapped MNase-seq reads^41^. Namely, we used a triweight kernel *K* as a weighting function (Equation 1) and the kernel-smoothed dyad counts D at each genomic coordinate *i* is calculated as ^40^ .

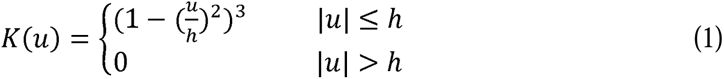

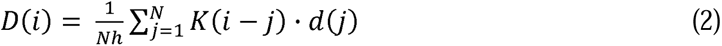

Here *N* is the length of a chromosome, *d(j)* is the dyad count at genomic coordinate *j* and *D(i)* is the smoothed dyad count at genomic coordinate *i*. Small values of bandwidth *h* lead to less smoothing but more accurate estimates of dyad positions. In our case, we chose a relatively small bandwidth value *h*=15 to improve the accuracy of the mapped dyad positions.

Next, we identified the genomic locations with the local maximum values of the smoothed dyad counts using bwtool^42^ and the minimum distance between the neighboring local maxima was set to 150 base pairs with “find local-extrema -maxima -min-sep=150”. Then, for every 60-bp window centered at each local maxima, one representative dyad location was determined as the dyad location with the highest number of dyad counts in this interval. Other dyad positions within the same 60-bp interval were discarded. If two or more dyad positions had the same dyad counts within the same interval, the dyad position located closest to the local maximum of the smoothed counts was selected as the representative dyad. Last, nucleosome regions (NRs) were determined as genomic regions centered at the representative dyad positions and flanked by 73 bp segments on each side.

### Genome-wide mapping of active enhancers, open chromatin and nucleosome-depleted regions

To map the open chromatin and enhancer regions, we used DNase-seq and H3K27ac and H3K4me1 ChIP-seq data from the following five human cell lines (H1, HepG2, MCF7, K562, and HeLa-S3, Supplementary Table 1 and 2). The narrowPeak files were downloaded from the ENCODE and Roadmap project^43,44^. The open chromatin regions were identified as genomic regions centered at narrow peaks and flanked by 1000 bp segments on each side. The active enhancer regions were defined as the open chromatin regions overlapped with both H3K27ac and H3K4me1 ChIP-seq narrow peaks. The nucleosome-depleted regions (NDRs) represent genomic regions free of nucleosomes and were identified as genomic regions not covered by any mono-nucleosome fragments (120 to 180 base pairs length) in MNase-seq data in all replicate experiments.

Using open chromatin regions from the DNase-seq data, we identified *differentially* and *conserved open chromatin* regions using the “intersect” command from the BEDTools suite^39^. Conserved open chromatin regions represent open chromatin regions that are more than 80% shared between embryonic H1 and at least one other differentiated cell line used in this study^45^ (Supplementary Figure 1). Chromatin regions with differential accessibility (“*differentially open chromatin regions*”) are defined as those that have less than 20% overlap between open regions in H1 embryonic cell line and open regions in one of differentiated cell lines so that these regions are closed in one cell line type and open in another^45^ (Supplementary Figure 1). While predicting pioneer transcription factors, that are important during the differentiation of embryonic stem cells (the majority of factors in Test set 1 and all factors in Test set 2), we defined differentially open chromatin regions as those closed in H1 and open in differentiated cell lines. Vice versa, based on their functions, for seven factors from Test set 3, which act in reprogramming somatic cells into induced pluripotent stem cells, we defined the differentially open regions as those closed in differentiated cell lines and open in the H1 embryonic cell line. For motif enrichment analysis of transcription factors, we selected two sets of nucleosome regions (NRs) and nucleosome-depleted regions (NDRs): 1) NDRs located in the open chromatin regions and all identified NRs using the MNase-seq data; 2) NDRs located in conserved open chromatin regions and NRs located in the *differentially* open regions so that their accessibility may be associated with TF binding.

### Analysis of dinucleotide patterns of nucleosomal DNA

Using identified representative nucleosome dyad positions, we examined dinucleotide patterns of nucleosomal DNA and mapped genomic coordinates of WW/SS (where W is A or T, and S is G or C) and YY/RR (R = A or G, and Y = C or T) dinucleotides on NRs. Then, we aligned NRs by superimposing their dyad positions and computed the frequency of observed dinucleotides at each location of nucleosomal DNA (as a function of distance in base pairs from the nucleosome dyad). As a result, we observed pronounced dinucleotide patterns at specific nucleosomal DNA positions for all cell lines which are indicative of the high qulity of the data and nucleosome mapping procedure (Supplementary Figure 2).

### Genome-wide mapping of transcription factor binding sites

To map the genome-wide locations of binding sites of various transcription factors, we matched the ChIP-seq data from the ENCODE project^43^ with the MNase-seq data for the same human cell lines (H1, HepG2, MCF-7, HeLa-S3 and K562) (Supplementary Table 3). In case of HeLa cell line, we used ChiP-seq data in HeLa-S3, which is a clonal derivative of HeLa. In total, ChIP-seq data for 225 transcription factors could be matched with the corresponding MNase-seq data from the same cell type. All available narrow peak files of ChIP-seq of these transcription factors were downloaded, and files corresponding to the same transcription factor were merged for further analysis. To map binding sites from ChIP-seq narrow peaks, referred to as “ChiP-seq TF motif”, we downloaded position frequency matrices (PFMs) for each TF from the JASPAR CORE database ^46^. Then we applied FIMO program ^47^ to scan the DNA sequences within ChIP-seq narrow peaks using each PFM and identified motifs with p-value less than 10^-4^ for further analysis.

Next, we calculated the transcription factor binding motif profiles or briefly *“motif profiles”*. To do that, we first mapped ChIP-seq TF motifs for a given TF to the closest nucleosome dyad position and then aligned all dyad positions and corresponding ChIP-seq TF motifs. For each TF binding motif, we considered sequences in both strands of DNA. Then, we counted the number of base pairs from different TF binding motifs that mapped at each location of nucleosomal DNA and flanking DNA regions (+/- 1000 bp) around the dyad.

### Transcription factor motif enrichment analysis

Pioneer transcription factors (PTFs) can engage nucleosomal DNA, while truly canonical TFs cannot bind to nucleosomes. To predict PTFs, we calculated the binding motif enrichment of different TFs on NRs compared to NDRs. Namely, we counted a number of base pairs of ChIP-seq TF motifs overlapped with the NR and NDR regions and constructed the following contingency table (Supplementary Table 4). We then calculated the enrichment score for each transcription factor (Equation 4) and applied a Fisher’s exact test to evaluate the significance of TF motif enrichment on NRs compared to NDRs.

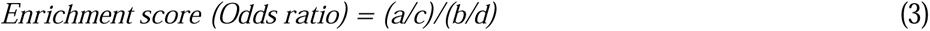

Here *a* and *c* are the number of base pairs overlapped with ChIP-seq TF motifs on NR and NDR respectively. Counts *b* and *d* correspond to the number of base pairs outside of ChIP-seq TF motifs on NR and NDR respectively. We further collected the data on expression levels of various TFs from the NIH roadmap epigenomics program (https://egg2.wustl.edu/roadmap/data/byDataType/rna/expression/) and identified TFs which are highly expressed in each cell line (RPKM value >= 10).

Receiver operating curves (ROC) and precision-recall (PR) curve analyses were further performed to evaluate the power of enrichment scores in discriminating pioneer transcription factors from other transcription factors. 32 known pioneer transcription factors from the literature (which were included in our list of 225 TFs) were used as positives (Supplementary Table 5) and other transcription factors were considered as negatives for this test.

### Clustering of transcription factor binding motif profiles

To identify the prevalent interaction modes of various TFs with nucleosomes, we have performed clustering of TF binding motif profiles. Prior to clustering, we filtered out potentially low-quality binding profiles using the following criteria. First, we removed under-represented TFs with the total genome-wide cumulative sum of ChiP-seq TF motifs on nucleosome regions of less than 500 base pairs. Second, due to the two-fold symmetry of DNA in nucleosome structures, ChiP-seq TF motifs on DNA complementary plus and minus strands should be structurally superimposed if the nucleosome structure is rotated by 180 degrees. If we are analyzing relative large number of binding motifs on both DNA strands, binding motif profiles should be symmetrical with respect to the nucleosome dyad. Therefore, we calculated the Pearson correlation coefficient of motif profiles between two symmetrical nucleosomal halves (positive and negative superhelical locations of nucleosomal DNA) for each TF and removed those with Pearson correlation coefficient values less than 0.4.

Next, we applied t-distributed stochastic neighbor embedding (t-SNE) and projected all profiles onto two dimensions using the Rtsne function from the R package^48^. Then, the projected data were subjected to k-medoids clustering using the pam function from the R package with the optimal number of clusters equal to six (Supplementary Figure 9). The silhouette width is an estimate of the goodness of clusters, its values close to 1 correspond to a cluster where most objects are much closer to other objects in the same cluster than to other clusters. For each cluster, members with silhouette width <= 0.25 were considered as outliers and 33 outliers removed.

Since Micrococcal Nuclease has a sequence bias and cleaves DNA upstream of A or T more efficiently than of G or C nucleotides, a certain nucleotide preference exists around the ends of nucleosomal DNA reads produced by the MNase-seq experiments. It may potentially bias the TF binding profiles near the nucleosomal DNA ends. Therefore, in our analysis, we excluded regions near the nucleosomal DNA ends. To identify binding modes between TFs and nucleosomes, we clustered binding motif profiles of different TFs centered at the nucleosomal dyad locations (+/- 60 base pair from dyad) using k-medoids.

## Results

### Transcription factor binding motif enrichment analysis can identify pioneer transcription factors

Nucleosomes are generally considered an impediment to the binding of TFs to DNA and thus binding sites of TFs are typically depleted in DNA regions with high nucleosome occupancy. However, pioneer TFs can recognize their binding motifs on nucleosomal DNA and trigger the opening of chromatin to recruit other TFs in a cell-type-specific fashion ^2,49^. Therefore, we hypothesized that DNA binding sites of pioneer transcription factors should not be depleted and, in some cases, should be enriched on nucleosome regions (NRs). At the same time, those canonical transcription factors that can preferentially bind to the naked DNA should exhibit the depletion of binding sites on nucleosome regions and enrichment in nucleosome-free regions. To quantitively assess these trends, we have performed binding motif enrichment analysis for each of the 225 transcription factors and calculated the motif enrichment scores (Supplementary Table 6-9). We have further analyzed the enrichment of TF binding sites on nucleosome regions (NRs) compared to the nucleosome depleted regions (NDRs)(see Methods for definition).

To validate our results we have compiled three sets of known pioneer transcription factors^2,12,14,16^(which could also be found in our list of 225 TFs) as positives (Supplementary Table 5 and 10), whereas other transcription factors were considered as negatives for this test. Test set 1 includes 32 known pioneer transcription factors from our data set. Some known PTFs were not included as they were not present in our original data sets. Test set 1 comprises Test set 2 and 3 and other known pioneer transcription factors. Test set 2 includes 11 known pioneer transcription factors with specific roles in cell differentiation. Test set 3 includes seven known pioneer transcription factors critical for the maintenance of embryonic stem cells or the reprogramming of somatic cells into induced pluripotent stem cells (Supplementary Table 10). As we show in the next section, the negative set may also contain pioneer transcription factors, therefore the classification accuracy values provided below can be considered as a lower bound estimates. Then, we performed the enrichment anlysis by calculating the binding motif enrichment of different TFs on nucleosomal regions compared to nucleosome-depleted regions.

The enrichment score of the 32 known PTFs from Test set 1 was found to be significantly higher than for other factors (p-value = 2.12*10^-7^ for all TFs and p-value = 4.33*10^-5^ for highly expressed TFs, Mann Whitney U test, Supplementary Figure 3 and Supplementary Table 11). The results also show the efficiency of enrichment scores for the classification of pioneer transcription factors (Supplementary Table 11, ROC AUC = 0.69, PR AUC = 0.33 and maximal MCC= 0.31 for all TFs and ROC AUC = 0.71, PR AUC = 0.39 and maximal MCC= 0.31 for highly expressed TFs).

Next the validation pertaining to the ability of pioneer transcription factors to open the closed chromatin was performed. We hypothesized that the enrichment score calculated based on NRs located in differentially open chromatin regions and NDRs located in conserved open chromatin regions would perform best in the classification of pioneer transcription factors essential for cell differentiation (Test set 2). As can be seen in Figure 2 and Supplementary Table 11, it is indeed the case and the classification accuracy increases from ROC AUC = 0.69 to 0.89 (from 0.71 to 92 for expressed TFs) upon the inclusion of differentially open regions compared to open regions (maximal Maximal Matthews correlation coefficient increased from 0.31 to 0.42). We found that known pioneer transcription factors that acted as key regulators of cell differentiation had the highest enrichment scores in our ranking (Figure 2c). These cases (Test set 2) mainly included pioneer transcription factors from the FOXA, GATA, and CEBP families ^50–52^ with GATA1 and GATA2 showing the highest enrichment scores.

**Figure 2.**
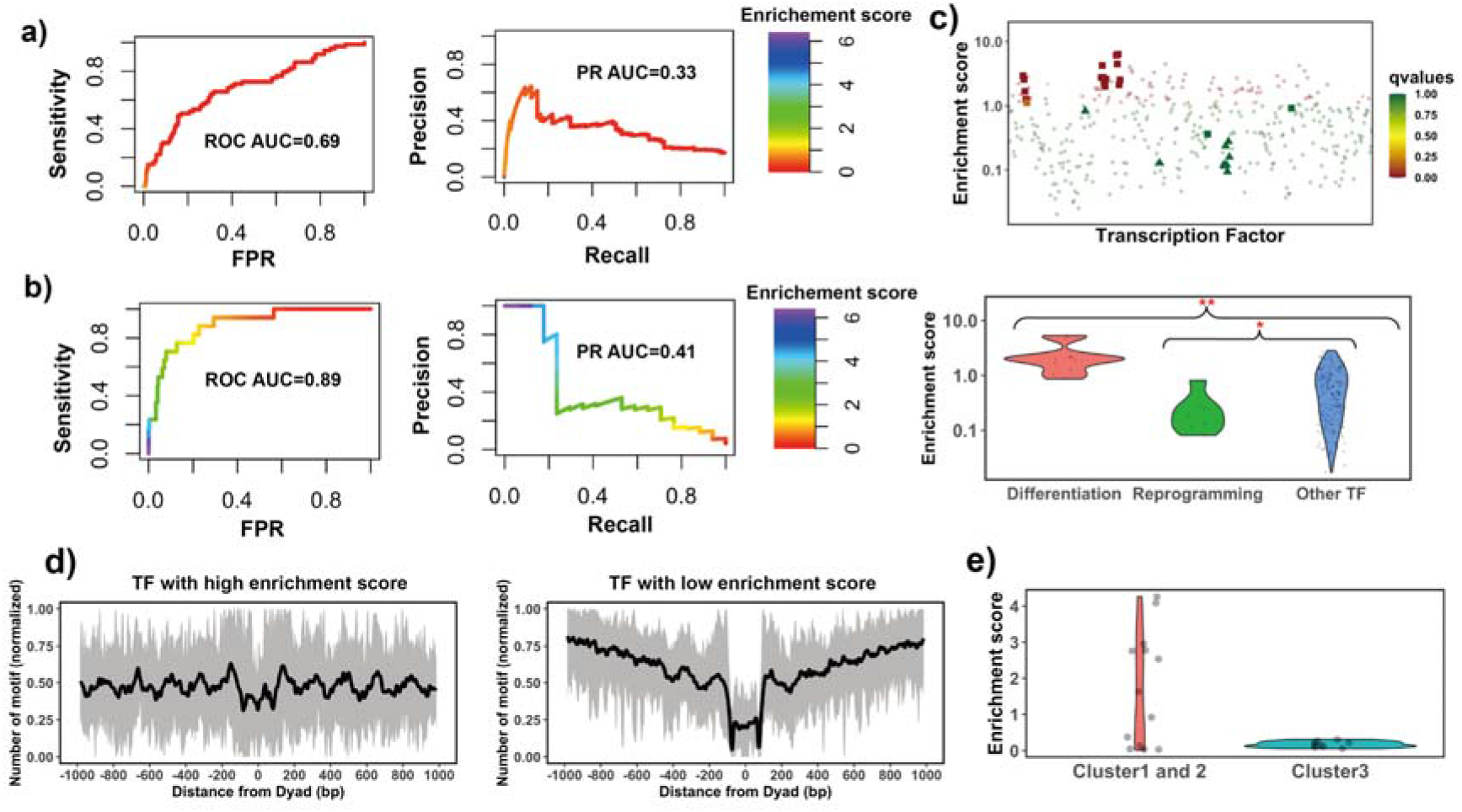
Identifying pioneer transcription factors using motif enrichment analysis. **a)** TF motif enrichment score is used to distinguish 32 known PTFs (Test set 1) from other other TFs. Receiver operating curve (ROC) and precision-recall (PR) curve analysis of motif enrichment scores are performed. Here nucleosome regions (NRs) were determined as genomic regions (147 bp long) centered at the representative dyad positions and the nucleosome-depleted regions (NDRs) represent genomic regions free of nucleosomes and are located in open chromatin regions. The random retrieval classifier would predict with AUC=0.5 and PR=the fraction of true positives=0.17. **b)** TF motif enrichment score is used to distinguish 11 known PTFs with essential roles in cell differentiation (Test set 2) from other other TFs. Here NRs in differentially open and NDRs in conserved open chromatin regions are used in enrichment analysis. The random retrieval classifier would predict with AUC=0.5 and PR= 0.04. **c)** Classification of pioneer transcription factors by binding motif enrichment scores. Known pioneer transcription factors from Test set 2 and Test set 3 are indicated by squares and triangles, while other TFs are shown as circles. Colors corresponds to false discovery rate (FDR) q-values. Mann–Whitney U tests are performed under the null hypothesis that PTF’s mean values of enrichment scores are equal to canonical TFs. * - p-value < 0.05; ** - p-value < 0.005. **d)** Binding motif profile of TFs with the highest and lowest motif enrichment scores (ranked at the top or bottom 20% among all TFs). The number of motifs for each TF is normalized within the range between 0 and 1 as follows: X(i)_normalized_) = (X(i) -X_min_)/(X_max_ -X_min_), X(i) is the number of sequences which have TF binding sites at the *i*_th_ base pair from the nucleosomal dyad position; X_max_ and X_min_ represent the maximal and minimal counts of sequence fragments. **e)** Comparison of the enrichment score of TFs in different clusters identified from recent EMSA experiments^19^. Only 13 transcription factors could be found in EMSA and our data set. Cluster 1 and 2 include strong binders to both naked DNA and nucleosomal DNA and weak binders to both naked DNA and nucleosomal DNA (only one TF). Cluster 3: strong binders to naked DNA but weak binders to nucleosomal DNA.

Interestingly, pioneer transcription factors in Test set 3 (responsible for the maintenance of embryonic stem cell or reprogramming of somatic cells) showed significantly lower enrichment scores compared to other TFs (Figure 2c and Supplementary Figure 4). Yamanaka pioneer transcription factors (POU5F1/OCT4 and KLF4)^53^ were strongly depleted at nucleosomes (Figure 2c). It has been previously shown that Yamanaka pioneer transcription factors might recognise partial sequence motifs on nucleosomal DNA and require other factors for their binding to nucleosomes^16^ and therefore their enrichment score might not be expected to be high. We also found known pioneer transcription factors with relatively low enrichment scores including NFYA, NFYB, NFYC and ESRRB. These, transcription factors regulate stem cell proliferation and maintenance of stem cell identity^54–56^ (Supplementary Figure 4).

However, when we repeated our analysis by redefining differentially open regions as those closed in differentiated cell lines and open in H1 embryonic cell line, then ESSRB and Yamanaka pioneer transcription factor POU5F1 (OCT4) showed significantly higher enrichment scores (Supplementary Figure 5). This could be explained by the roles of Yamanaka factors in cellular reprogramming – they reprogram somatic differentiated cells into induced pluripotent stem cells.

Since PTFs often target enhancers, we have repeated the enrichment anlysis using NRs located in differentially active enhancer regions and NDRs located in the conserved active enhancer regions and the performance in the PTF classification was slightly worse (Supplementary Figure 6). Here, differential enhancer regions refer to the active enhancer regions in an embryonic cell line (H1) that have less than 20% overlap with any active enhancer regions in differentiated cell lines. Conserved active enhancer regions represent active enhancer regions that are more than 80% shared between embryonic H1 and at least one other differentiated cell line used in this study. We also explored different thresholds in defining *differentially open* and *conserved open chromatin regions* but the performance in classifications of PTF was not significantly affected by the threshold choice (Supplementary Figure 7).

Finally, we performed an additional validation using recent data from the high-throughput protein microarray and electromobility shift assays (EMSA) experiments on human TFs which systematically assessed transcription factor binding preferences to nucleosomal DNA versus naked DNA^19^. These authors classified all studied transcription factors with respect to their strengths of nucleosome binding into three clusters: strong binders, which bind strongly to both naked DNA and nucleosomal DNA (cluster 1) and weak binders, which bind weakly to both naked DNA and nucleosomal DNA (cluster 2). Cluster 3 consists of strong binders which bind strongly to naked DNA but weakly to nucleosomal DNA. We found that TFs from their cluster 3 had the lowest enrichment scores although this trend was not statistically significant because of a lack of the data in this cluster. As to TFs in cluster 1 and 2, half of them had binding sites enriched on nucleosome regions (including known pioneer transcription factors FOXA1, GATA4 and CEBPA) which correspond to strong nucleosome binders, and another half had binding sites enriched on nucleosome depleted regions (Figure 2e).

Using gene expression data from the Roadmap Epigenomics Program^57^ and enrichment analysis on all open chromatin regions, we have identified 35 TFs in H1, K562, HepG2 and HeLa cell lines (no expression data was available for MCF-7) that ranked highly in the enrichment analysis and were also highly expressed in the corresponding cell lines (Supplementary Table 7). Among these 35 TFs, 15 TFs were well-characterized PTFs such as GATA, FOXA and CEBP factors, ESRRB, NEUROD1, SPI1 and subunits of the AP-1 complex. To validate the remaining 20 PTF predictions, we performed literature searches and found that RFX5 had the ability to displace nucleosomes^14^. In addition, many of the predicted PTFs such as ZKSCAN1, USF1,USF2 and SRF, were confirmed by a recent study^58^.

Next, using the enrichment score calculated based on NRs located in differentially open chromatin regions and NDRs located in conserved open chromatin regions, we identified 40 transcription factors that could act as PTFs with essential roles in cell differentiation in K562, HepG2 and HeLa cell lines. These identified TFs had their DNA binding sites significantly enriched on nucleosomal DNA of differentially open chromatin regions and were highly expressed in the corresponding cell lines (Supplementary Table 9). Among these 40 TFs, 15 were well-characterized PTFs including GATA, FOXA and CEBP factors and subunits of AP-1 complex. For the remaining 25 PTF predictions, seven transcription factors were annotated in the literature as potential pioneer transcription factors and/or potential nucleosome binders (Supplementary Table 12). For instance, HNF4A was annotated as a potential pioneer transcription factor active in chromatin remodeling in the liver^59^, LEF1 was identified as a regulatory high mobility group (HMG) box protein that could bind to nucleosomes ^60^ and CUX1 could specifically interact with its recognition motif in a nucleosomal context ^61^.

### Association between binding motif profiles and nucleosome occupancy for pioneer transcription factors

The enrichment analysis described above is the first step to estimate the propensity of transcription factor binding sites to be located on nucleosomal footprints. The enrichment analysis might have high specificity but a compromised sensititivity as it can misclassify those pioneer transcription factors that can bind to both naked DNA and nucleosomes at similar concentrations. Therefore, the next step would be to evaluate the actual locations of TF binding sites with respect to the nucleosomal dyad. To this end, we tested if there is a significant association between the binding motif profiles (see Methods) and nucleosome occupancy values +/-400 bp around the dyad (Figure 3a). Our results showed that 87% of all 225 transcription factors had negative PCC between binding motif profile and nucleosome occupancy values (Figure 3b) which is consistent with the fact that nucleosomes generally restrict the access of TFs to their binding sites on DNA molecules.

**Figure 3.**
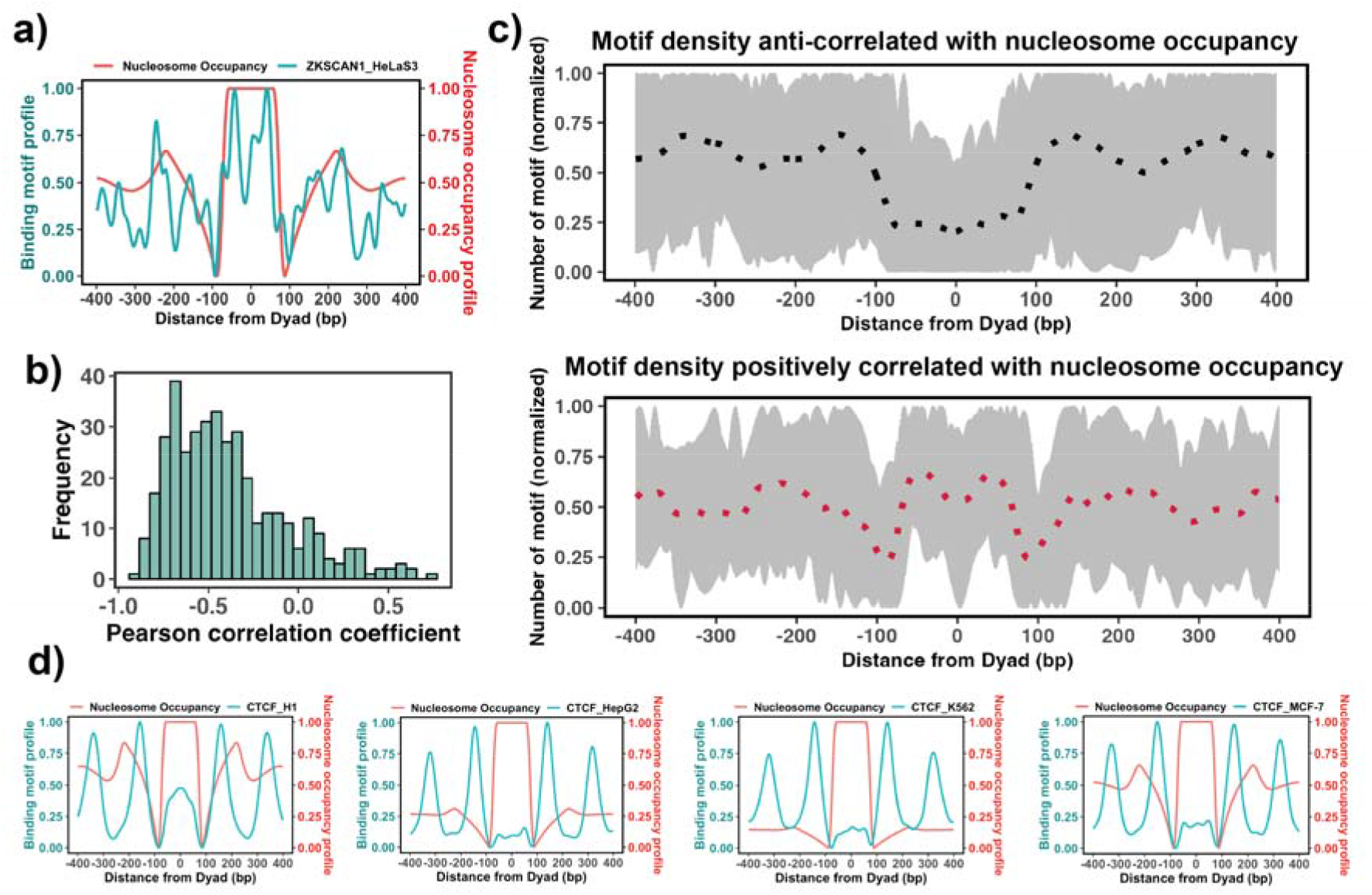
Association between the binding motif profiles and nucleosome occupancy. **a)** Motif profiles for ZKSCAN1 in HeLa-S3 cell line is shown as an example. **b)** Pearson correlation coefficients between motif profiles and nucleosome occupancy values for each TF (n=225) (the median value of correlation coefficient = - 0.46). **c)** Binding motif profiles of TFs with positive (red, correlation coefficient >= 0.2 and p-value < 0.05) or negative correlation coefficients (black, correlation coefficient <= -0.4 and p-value < 0.05) between motif profile and nucleosome occupancy. Dashed lines correspond to the average of binding motif profiles. **d)** Comparison of binding motif profiles of CTCF between H1 embryonic stem cell line and somatic cell lines.

We have also identified 37 transcription factors that had a statistically significant positive correlation coefficients between binding motif profile and nucleosome occupancy and could be classified as potential nucleosome binders and pioneer transcription factors (Supplementary Table 13 and Figure 3). Among these predictions there were six known PTFs, such as POU5F1(OCT4), GATA3, CEBPB, ATF2, NFYA and NFYB (Supplementary Table 13). As we mentioned previously, NFYB, NFYA and POU5F1 had low enrichment scores, but were identified as nucleosome binders by this correlation analysis. It could be explained by their dual nature: these factors can bind to nucleosomes using certain binding arrangements, as evident from their postive correlation coefficients, but at the same time, they can interact with the naked DNA, as evident from their low enrichment scores. Supplementary Table 13 shows 14 predictions that overalpped with the predicted PTFs from the enrichment analysis (ATF2, BACH1, CEBPB, ESRRA, HMBOX1, NFATC3, NFYB, SREBF2, USF1, USF2, ZNF24, ZNF274, ZNF282, ZKSCAN1). Among those with a high PCC > 0.5, were ZKSCAN1, ESR1, NFATC3, ZBTB7B, MAX, and TBX2. Half of them were also predicted to be pioneers by a recent study^58^. With the exception of a few cases, none of transcription factors was identified as being pioneer in all cell types because pioneer activity is often cell-type specific. For example, CTCF was identified as having a significant correlation (although low) between binding motif profile and nucleosome occupancy for the human embryonic stem cell line (PCC = 0.1, p-value < 0.05), but not for other somatic cell lines (Figure 3d). Indeed, previous studies have indicated that CTCF proteins could access the binding sites in nucleosomes and may function as pioneer transcription factors in embryonic stem cells and to a lesser extent in differentiated cells^62–64^.

### Deciphering the interaction modes between transcription factors and nucleosomes

A recent study characterized the interaction landscape between pioneer transcription factors and nucleosomes using NCAP–SELEX (Nucleosome Consecutive Affinity-Purification with Systematic Evolution of Ligands by Exponential Enrichment) ^14^. It revealed different binding modes of pioneer transcription factors: DNA end binding, dyad binding, gyre binding and periodic binding^14^. The NCAP–SELEX approach was based on the analysis of enrichment of specific sequences from the DNA libraries. These sequences were reconstituted into nucleosomes, and incubated with TFs^14^. Then, the dissociated nucleosomal DNA was separated from the intact nucleosomes and the analysis of the enrichment of sequences allowed to identify TF binding specificities and binding site locations on nucleosomal DNA^14^. To compare our TF motif profiles with NCAP–SELEX data, we filtered out low-quality motif profiles using the criteria described in the Methods section and then calculated motif enrichment scores for TFs with different binding modes identified by NCAP–SELEX approach. Due to the limited number of transcription factors observed in both our dataset and NCAP–SELEX study (24 transcription factors), we mainly focused on the DNA end binding super helical locations (SHL) from +/- 5.5 to +/- 7 and the dyad binding modes with SHL from 0 to +/- 1.5. To estimate the preferential binding of TFs to the ends of nucleosomal DNA compared to the nucleosomal dyad, we calculated the end/dyad binding ratio (R) as the number of binding motifs at the DNA ends (SHLs from +/-5.5 to +/-7) divided by the number of binding motifs near dyad regions (SHLs from 0 to +/-1.5).

In NCAP-SELEX experimental analyses, to quantify the preference of PTF binding to nucleosomes^14^, binding signals were compared by calculating the mutual information (MI) content between 3-mer distributions at two non-overlapping positions of the ligand, aimed at finding if SELEX ligand may contact these positions at the same time. Since nucleosomes can form on most DNA sequences, whereas TFs bind to only a few specific sequences, the NCAP-SELEX study calculated the enriched MI (E-MI) score to separate the TF signals from nucleosome signals by limiting the MI measure to the top ten most enriched 3-mer pairs. E-MI penetration score corresponds to E-MI drop by half compared to the E-MI maximum and larger values pointed to the favorable binding to the dyad regions^14^. As our R and experimental EMI penetration values should be anti-correlated, we indeed observed a statistically significant negative correlation between R_end/dyad_ and EMI penetration values (PCC = -0.45, p-value = 0.025). This shows that our computational analysis is able to capture the detailed features of TF binding motifs on nucleosomes. For the EMI intensity, although a negative linear dependence trend was evident, the correlation was not significant (Supplementary Figure 8).

### Clustering reveals several groups of pioneer transcription factor binding sites

Previous studies indicated that while nucleosome positioning acted as a barrier to TF binding, there was a number of different signatures of mutual TF/nucleosome positioning^65^. We attempt to characterize the details of PTF binding motifs at near-single nucleotide resolution on nucleosomal DNA and performed K-medoids clustering of binding motif profiles for all 225 transcription factors (Figure 4 and Supplementary Figure 9). Transcription factors in the first and second cluster showed that motif density increased with the distance from the dyad. These clusters include cases of canonical factors or those PTF that preferentially interact with the ends of nucleosomal DNA from super helical locations (SHL) +/- 5.5 to SHL +/- 6. Such pioneer transcription factors include CEBPB and GATA3, consistent with the NCAP-SELEX data. In addition, it has been recently shown that GATA3 factor targets nucleosomal DNA around the SHL +/-5.5 position^23^. In clusters 4 and 6, TFs preferentially interact with the nucleosomal DNA around the dyad regions and SHL from +/- 3 to SHL +/- 5, whereas transcription factors from clusters 3 and 5 preferentially occupy SHL +/- 1 to SHL +/- 2.5 respectively.

**Figure 4.**
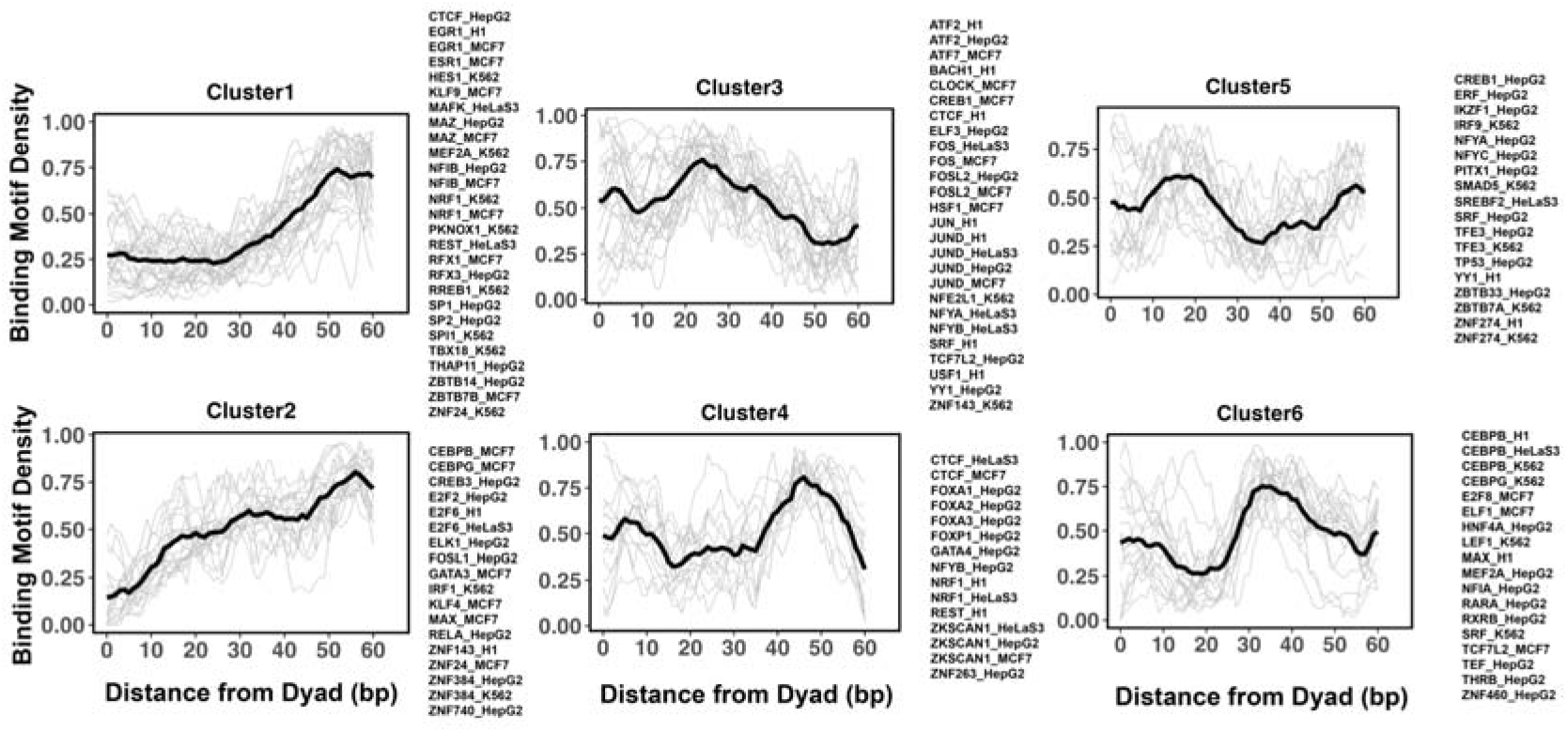
Clusters of TF binding motif profiles on nucleosomal DNA. Binding motif profiles centered at nucleosomal dyad locations (+/- 60 base pair from dyad) are clustered using k-medoids clustering with k=6. The entry/exit regions of nucleosomal DNA were excluded as a certain nucleotide bias exists around the ends of nucleosomal DNA reads produced by the MNase-seq experiments. Binding motif profiles between two symmetrical nucleosomal halves are combined for each TF. The black line represents the averaged profiles of all TFs in the same cluster. Cluster members with silhouette width <= 0.25 were considered as outliers and removed.

We found that many transcription factors from the same family and even from different cell types were assigned to the same cluster. For example, all known pioneer transcription factors from FOXA families belong to cluster 4 and indeed it has been shown that FOXA1 can target its binding motifs to nucleosomal DNA near the dyad and linker regions^66^. However, there are a few exceptions, CEPBP, FOSL, CREB1, CTCF, MEF2A, NFYA, NFYB and YY1, where the same transcription factor from different cell lines has been assigned to different clusters. Possible reasons may include binding motifs on DNA which can be close in space but occupy different regions on DNA. In addition, the cell-type-specific epigenetic marks recognized by pioneer transcription factors or cooperativity binding between factors are differentially expressed in various cell types.

## Discussion

The modulation of chromatin accessibility with high spatiotemporal precision is a subject of ongoing debate as it is a crucial factor in regulation of transcription, replication and DNA repair processes. The question of how chromatin accessibility is precisely modulated involves multiple aspects, including epigenetic modifications, PTF infiltration, ATP-dependent remodeling and spontaneous chromatin dynamics. There are about two thousand transcription factors in the human genome and a few hundred of them may have PTF properties. Yet, for the vast majority of human PTFs, the locations of their binding sites and mechanisms of binding and regulation remain unknown^19,67^. There are many different experimental assays that can provide information on transcription factor binding sites but these assays suffer from multiple drawbacks and often require prior knowledge of the TFs being tested. Moreover, pioneer transcription factors are very dynamic, may target partial binding sites and work cooperatively with other factors. For example, two recent cryo-EM structures of the pioneer transcription factor SOX2-nucleosome complexes showed different mechanisms of binding^21,22^. All this complicates the identification and characterization of human pioneer transcription factors pointing to the pressing need to develop predictors and classifiers ^49,68–70^ . The goal of this study has been to gain functional insights into the mechanisms of binding and infiltration of PTFs into chromatin at the level of the nucleosome. To achieve this goal, we used ChIP-seq, MNaseq-seq and DNase-seq data from five different cell lines for 225 human transcription factors. A computational framework has been to systemically investigate the ability of transcription factors to bind to nucleosomes. As a result, we found that using the information on differentially open chromatin regions (open in one cell line, closed in another) leads to the highest classification accuracy in discriminating pioneer from canonical transcription factors. This finding supports the view that TF binding to nucleosomes leads to DNA and chromatin opening and might correlate with the reprogramming potential^19^. Our study has also verified the known and predicted several dozens of new PTFs as nucleosome binders (Supplementary Tables 7, 9 and 13). These predicted cases include TFs without previously known pioneer activity which could be subject to the future experimental validation. Finally, we have identified six distinctive clusters of TF binding profiles with the nucleosomal DNA. These clusters point to the diversity of binding motifs where transcription factors belonging to the same cluster may exhibit potential competitive binding.

We should mention that our classification method and the data used in this study have certain limitations. First, interactions of PTFs with nucleosomes may depend on binding of other factors. Second, PTF binding relies on specific recognition of DNA binding sites and can be governed by nucleosome dynamics. Indeed, initial stages of binding can happen via the DNA site exposure through thermal fluctuations, leading to DNA unwrapping and the formation of DNA looping or twist defects^71–73^. Pioneer transcription factors can exploit DNA unwrapping and trap nucleosomes in a partially unwrapped states^74^. Transcription factor binding motifs close to the nucleosome entry-exit sites may have increased exposure but would be error-prone to capture using MNase-seq data as a nucleotide bias exists around the ends of nucleosomal DNA reads produced by the MNase-seq experiments. Moreover, as we showed recently, the degree of spontaneous DNA unwrapping from nucleosomes depends on the incorporation of histone variants, like H2A.Z^75^, which is, in turn, might be needed for pioneer transcription factor recruitment to promote embryonic stem cell differentiation^76^. The third group of limitations pertains to the nucleosome positioning in a genome, as binding dissociation constants for PTFs depend on where nucleosomes are located. Indeed, nucleosome positioning and stability is not uniform throughout the genome and depends on local nucleosomal DNA sequence, histone variant deposition and epigenetic chromatin modifications. Finally, pioneer transcription factors may exhibit multivalent binding recognizing not only DNA binding sites but also some parts of the histone core or histone tail regions. Deducing such dependencies from the experimental assays utilized in this study is very challenging, if not impossible.

Nucleosomes represent hub points in epigenetic signaling pathways and identifying complex epigenetic relationships at the level of single nucleosomes may yield functional insights into the mechanisms of binding and infiltration of pioneer transcription factors into chromatin during differentiation and reprogramming. Transcription factors regulate a large number of signaling pathways and their dysregulation contributes to a plethora of human diseases, including diabetes, cardiovascular diseases and many cancers. Conventional transcription factors have been used for a long time as biomarkers and drug targets, however, the targeted therapeutic potential of pioneer transcription factors is lagging. To fill this gap, integrative approaches using large-scale low- or medium-resolution data with precise molecular modeling or protein-protein docking may provide the required detailed characterization of many predicted PTFs in the future.

## Acknowledgments

ARP was supported by the Department of Pathology and Molecular Medicine, Queen’s University, Canada. ARP is the recipient of a Senior Canada Research Chair in Computational Biology and Biophysics and a Senior Investigator Award from the Ontario Institute of Cancer Research, Canada. ARP acknowledges the support of the Natural Sciences and Engineering Research Council of Canada (NSERC) (No. RGPIN/02972-2021 ARP). ARP would like to thank David Clark for helpful discussions. YP was supported by the National Natural Science Foundation of China (No.12205112) and Fundamental Research Funds for Central China Normal University. YP, WS, IO and DL were supported by the Intramural Research Program of the National Library of Medicine, NIH. VBT acknowledges support by Cancer Research UK (grants EDDPMA-Nov21\100044 and SEBPCTA-2022/100001) and BBSRC IAA grant.

