## Supplemental Information for "Detection of new pioneer transcription factors as cell-type specific nucleosome binders"

**Supplementary Table 1**. Summary of MNase-seq experimental data for genome-wide nucleosome mapping.

| Cell lines | Sequencing type | GEO Accession number |
| --- | --- | --- |
| MCF-7 breast cancer cell line | paired-end | GSE51097 |
| K562 immortalized myelogenous leukemia cell line | paired-end | GSE78984 |
| H1 human embryonic stem cell line | paired-end | GSM1194220 |
| HepG2 human liver cancer cell line | paired-end | GSM3718063 |
| HeLa immortalized cervical tumor cell line | paired-end | GSE100401 |

**Supplementary Table 2**. Summary of the number of identified nucleosome regions (NRs) in each cell line using the reads with fragment sizes between 146-148.

| Cell lines | number of NRs |
| --- | --- |
| MCF-7 breast cancer cell line | 10,290,828 |
| K562 immortalized myelogenous leukemia cell line | 2,780,423 |
| H1 human embryonic stem cell line | 12,551,429 |
| HepG2 human liver cancer cell line | 5,193,194 |
| HeLa immortalized cervical tumor cell line | 9,842,040 |

**Supplementary Table 3**. List of the analyzed transcription factors and corresponding identifiers of position frequency matrices (PFMs) files from the JASPER database.

| TF gene name | Motif ID | TF gene name | Motif ID | TF gene name | Motif ID | TF gene name | Motif ID |
| --- | --- | --- | --- | --- | --- | --- | --- |
| ATF2 | MA1632.1 | **GABPA** | MA0062.3 | **MYBL2** | MA0777.1 | **SPDEF** | MA0686.1 |
| ATF3 | MA0605.2 | **GATA1** | MA0035.4 | **MYC** | MA0147.3 | **SPI1** | MA0080.5 |
| ATF4 | MA0833.2 | **GATA2** | MA0036.3 | **NEUROD1** | MA1109.1 | **SREBF1** | MA0595.1 |
| ATF6 | MA1466.1 | **GATA3** | MA0037.3 | **NFATC3** | MA0625.1 | **SREBF2** | MA0596.1 |
| ATF7 | MA0834.1 | **GATA4** | MA0482.2 | **NFE2** | MA0841.1 | **SRF** | MA0083.3 |
| BACH1 | MA1633.1 | **GFI1** | MA0038.2 | **NFE2L1** | MA0089.2 | **STAT3** | MA0144.2 |
| BCL6 | MA0463.2 | **GMEB2** | MA0862.1 | **NFE2L2** | MA0150.1 | **TBX18** | MA1565.1 |
| BHLHE40 | MA0464.2 | **HES1** | MA1099.2 | **NFIA** | MA0670.1 | **TBX2** | MA0688.1 |
| CEBPA | MA0102.4 | **HEY1** | MA0823.1 | **NFIB** | MA1643.1 | **TBX3** | MA1566.1 |
| CEBPB | MA0466.2 | **HINFP** | MA0131.2 | **NFIC** | MA0161.2 | **TCF3** | MA0522.3 |
| CEBPD | MA0836.2 | **HLF** | MA0043.3 | **NFIL3** | MA0025.2 | **TCF7** | MA0769.2 |
| CEBPG | MA0838.1 | **HMBOX1** | MA0895.1 | **NFIX** | MA0671.1 | **TCF7L2** | MA0523.1 |
| CLOCK | MA0819.1 | **HNF1A** | MA0046.2 | **NFKB2** | MA0778.1 | **TCFL5** | MA0632.2 |
| CREB1 | MA0018.4 | **HNF1B** | MA0153.2 | **NFYA** | MA0060.3 | **TEAD1** | MA0090.3 |
| CREB3 | MA0638.1 | **HNF4A** | MA0114.4 | **NFYB** | MA0502.2 | **TEAD2** | MA1121.1 |
| CREB3L1 | MA0839.1 | **HNF4G** | MA0484.2 | **NFYC** | MA1644.1 | **TEAD4** | MA0809.2 |
| CREM | MA0609.2 | **HOXA5** | MA0158.2 | **NKX3-1** | MA0124.1 | **TEF** | MA0843.1 |
| CTCF | MA0139.1 | **HSF1** | MA0486.2 | **NR2C1** | MA1535.1 | **TFAP4** | MA0691.1 |
| CTCFL | MA1102.2 | **HSF2** | MA0770.1 | **NR2C2** | MA0504.1 | **TFCP2** | MA0145.3 |
| CUX1 | MA0754.1 | **IKZF1** | MA1508.1 | **NR2F1** | MA0017.2 | **TFDP1** | MA1122.1 |
| DBP | MA0639.1 | **IRF1** | MA0050.2 | **NR2F2** | MA1111.1 | **TFE3** | MA0831.2 |
| DLX6 | MA0882.1 | **IRF2** | MA0051.1 | **NR3C1** | MA0113.3 | **TF_name** | Motif_ID |
| E2F1 | MA0024.3 | **IRF3** | MA1418.1 | **NR4A1** | MA1112.2 | **TGIF2** | MA0797.1 |
| E2F2 | MA0864.2 | **IRF5** | MA1420.1 | **NR5A1** | MA1540.1 | **THAP1** | MA0597.1 |
| E2F3 | MA0469.3 | **IRF9** | MA0653.1 | **NRF1** | MA0506.1 | **THAP11** | MA1573.1 |
| E2F4 | MA0470.2 | **ISL2** | MA0914.1 | **NRL** | MA0842.2 | **THRB** | MA1574.1 |
| E2F6 | MA0471.2 | **ISX** | MA0654.1 | **ONECUT1** | MA0679.2 | **TP53** | MA0106.3 |
| E2F7 | MA0758.1 | **JUN** | MA0488.1 | **ONECUT2** | MA0756.1 | **USF1** | MA0093.3 |
| E2F8 | MA0865.1 | **JUNB** | MA0490.2 | **OVOL1** | MA1544.1 | **USF2** | MA0526.3 |
| EGR1 | MA0162.4 | **JUND** | MA0491.2 | **PBX2** | MA1113.2 | **VEZF1** | MA1578.1 |
| ELF1 | MA0473.3 | **KLF10** | MA1511.1 | **PITX1** | MA0682.2 | **XBP1** | MA0844.1 |
| ELF2 | MA1483.1 | **KLF11** | MA1512.1 | **PKNOX1** | MA0782.2 | **YY1** | MA0095.2 |
| ELF3 | MA0640.2 | **KLF13** | MA0657.1 | **POU5F1** | MA1115.1 | **ZBED1** | MA0749.1 |
| ELF4 | MA0641.1 | **KLF16** | MA0741.1 | **PPARG** | MA0066.1 | **ZBTB12** | MA1649.1 |
| ELK1 | MA0028.2 | **KLF4** | MA0039.4 | **PRDM1** | MA0508.3 | **ZBTB14** | MA1650.1 |
| ELK3 | MA0759.1 | **KLF6** | MA1517.1 | **PRDM4** | MA1647.1 | **ZBTB26** | MA1579.1 |
| ELK4 | MA0076.2 | **KLF9** | MA1107.2 | **RARA** | MA0729.1 | **ZBTB33** | MA0527.1 |
| ERF | MA0760.1 | **LBX2** | MA0699.1 | **RBPJ** | MA1116.1 | **ZBTB7A** | MA0750.2 |
| ESR1 | MA0112.3 | **LEF1** | MA0768.1 | **RELA** | MA0107.1 | **ZBTB7B** | MA0694.1 |
| ESRRA | MA0592.3 | **MAFF** | MA0495.3 | **REST** | MA0138.2 | **ZEB1** | MA0103.3 |
| ESRRB | MA0141.3 | **MAFG** | MA0659.2 | **RFX1** | MA0509.2 | **ZKSCAN1** | MA1585.1 |
| ETS1 | MA0098.3 | **MAFK** | MA0496.3 | **RFX3** | MA0798.2 | **ZKSCAN5** | MA1652.1 |
| ETS2 | MA1484.1 | **MAX** | MA0058.3 | **RFX5** | MA0510.2 | **ZNF143** | MA0088.2 |
| ETV1 | MA0761.2 | **MAZ** | MA1522.1 | **RFX7** | MA1554.1 | **ZNF148** | MA1653.1 |
| ETV4 | MA0764.2 | **MEF2A** | MA0052.4 | **RORA** | MA0071.1 | **ZNF24** | MA1124.1 |
| ETV5 | MA0765.2 | **MEF2D** | MA0773.1 | **RREB1** | MA0073.1 | **ZNF263** | MA0528.2 |
| ETV6 | MA0645.1 | **MEIS1** | MA0498.2 | **RUNX1** | MA0002.1 | **ZNF274** | MA1592.1 |
| FOS | MA0476.1 | **MEIS2** | MA0774.1 | **RXRB** | MA0855.1 | **ZNF282** | MA1154.1 |
| FOSL1 | MA0477.2 | **MGA** | MA0801.1 | **SIX1** | MA1118.1 | **ZNF317** | MA1593.1 |
| FOSL2 | MA0478.1 | **MITF** | MA0620.3 | **SMAD3** | MA0795.1 | **ZNF382** | MA1594.1 |
| FOXA1 | MA0148.4 | **MIXL1** | MA0662.1 | **SMAD5** | MA1557.1 | **ZNF384** | MA1125.1 |
| FOXA2 | MA0047.3 | **MLX** | MA0663.1 | **SNAI1** | MA1558.1 | **ZNF460** | MA1596.1 |
| FOXA3 | MA1683.1 | **MNT** | MA0825.1 | **SOX13** | MA1120.1 | **ZNF652** | MA1657.1 |
| FOXK1 | MA0852.2 | **MNX1** | MA0707.1 | **SOX18** | MA1563.1 | **ZNF740** | MA0753.2 |
| FOXK2 | MA1103.2 | **MSX2** | MA0708.1 | **SP1** | MA0079.4 | **ZSCAN29** | MA1602.1 |
| FOXO4 | MA0848.1 | **MTF1** | MA0863.1 | **SP2** | MA0516.2 |  |  |
| FOXP1 | MA0481.3 | **MXI1** | MA1108.2 | **SP4** | MA0685.1 |  |  |

**Supplementary Table 4.** A contingency table for binding enrichment analysis to quantify the TF’s binding preferences on NRs or NDRs. Numbers are given in base pairs.

|  | **ChIP-seq TF motifs** | **Outside of ChIP-seq TF motifs** |
| --- | --- | --- |
| **Nucleosome Regions** | a | b |
| **Nucleosome Depleted Regions** | c | d |

**Supplementary Table 5.** A list of well-characterized known pioneer transcription factors (PTFs) from multiple literature studies.

| Transcription Factor Name | Function |
| --- | --- |
| FOXA1, FOXA2, FOXA3 | Tissue-specific gene activation; embryonic development, establishment of tissue-specific gene expression and regulation of gene expression in differentiated tissues. |
| GATA1, GATA2, GATA3, GATA4 | Early stages of cell differentiation and organ development across a variety of tissues; development and maintenance of hematopoietic systems. |
| CEPBA, CEBPB | Fate decisions during myeloid differentiation; essential for maintaining homeostasis of both embryonic and adult tissues; key regulators of hepatocyte differentiation. |
| NFY complex: NFYA, NFYB, NFYC | NF-Y, also known as the CCAAT-binding factor CBF, is a ubiquitously expressed heterotrimeric TF composed of NF-YA, NF-YB, and NF-YC subunits. Cell type-specific master transcription factors; NF-Y complex is required for the maintenance of embryonic stem cell (ESC) identity and is an essential component of the core pluripotency network. |
| ESRRB | Suppress cell differentiation and sustain ESC self-renewal |
| POU5F1 (OCT-4) | Yamanaka transcription factors  cellular reprogramming of somatic cells into induced pluripotent stem cells. |
| KLF4 |  |
| NEUROD1 | Neuronal reprogramming/neuronal conversion from human fibroblasts; neuronal differentiation. |
| TP53 | DNA repair, cell-cycle arrest, and apoptosis; tumor suppressor. |
| AP-1 complex: FOS, FOSL, FOSL2, MAFG, MAFF, MAFK, JUN, JUNB, JUND, ATF2, ATF3, ATF4, ATF6, ATF7 | The transcription factor AP-1 is a heterodimeric protein, composed of members of the basic region leucine zipper protein superfamily, specifically, the Jun, Fos, and activating transcription factor proteins, which regulate gene expression in response to a variety of stimuli, including cytokines, growth factors, stress, and bacterial and viral infections. |
| SPI1(PU.1) | PU.1 is an ETS-family transcription factor that plays a broad range of roles in hematopoiesis. A direct regulator of myeloid, dendritic-cell, and B cell functional programs, and a well-known antagonist of terminal erythroid cell differentiation. |

**Supplementary Table 6.** Top 25% transcription factors found by the enrichment analysis by calculating the binding motif enrichment on nucleosomal regions (NRs) compared to nucleosome-depleted regions (NDRs).

| TF_name | Cell line | Enrichment score | TF_name | Cell line | Enrichment score |
| --- | --- | --- | --- | --- | --- |
| ATF2 | HepG2, K562 | 0.65, 0.43 | **MNT** | HepG2 | 0.25 |
| ATF3 | K562 | 0.22 | **NEUROD1** | MCF-7 | 0.15 |
| ATF4 | K562 | 0.46 | **NFATC3** | K562 | 0.14 |
| ATF7 | HepG2, MCF-7 | 0.59, 0.37 | **NFE2** | K562 | 0.18 |
| BCL6 | K562 | 0.16 | **NFE2L1** | K562 | 0.13 |
| CEBPB | K562, HepG2, HeLa-S3, MCF-7, H1 | 0.47, 0.26, 0.25, 0.24, 0.19 | **NFE2L2** | HepG2, HeLa-S3 | 0.21, 0.13 |
| CEBPG | K562, MCF-7 | 0.52, 0.37 | **NFIB** | MCF-7 | 0.22 |
| CREB1 | HepG2, MCF-7 | 0.26, 0.25 | **NFIC** | K562 | 0.13 |
| CTCF | HeLa-S3 | 0.13 | **NFIX** | K562 | 0.31 |
| CUX1 | MCF-7, K562 | 0.38, 0.23 | **NFYB** | HeLa-S3 | 0.14 |
| ESR1 | MCF-7 | 0.19 | **NR2C2** | HeLa-S3 | 0.15 |
| ESRRB | K562 | 0.14 | **NR2F2** | MCF-7 | 0.15 |
| FOS | MCF-7, HeLa-S3 | 0.20, 0.15 | **ONECUT1** | HepG2 | 0.19 |
| FOSL1 | K562 | 0.14 | **PBX2** | K562 | 0.27 |
| FOXA1 | MCF-7, K562 | 0.21, 0.16 | **PKNOX1** | K562, MCF-7 | 0.13, 0.13 |
| FOXA2 | HepG2 | 0.15 | **RBPJ** | K562 | 0.16 |
| FOXA3 | K562 | 0.20 | **REST** | HeLa-S3, H1 | 0.26, 0.14 |
| FOXO4 | K562 | 0.19 | **RFX1** | MCF-7, K562 | 0.20, 0.18 |
| GATA3 | MCF-7 | 0.20 | **RFX5** | HeLa-S3 | 0.18 |
| HLF | HepG2 | 0.31 | **SPI1** | K562 | 0.14 |
| HMBOX1 | K562 | 0.24 | **SREBF1** | MCF-7 | 0.23 |
| IKZF1 | HepG2 | 0.97 | **SREBF2** | HeLa-S3 | 0.15 |
| IRF1 | K562 | 0.19 | **SRF** | H1, K562, MCF-7, HepG2 | 0.30, 0.20, 0.15, 0.14 |
| IRF2 | K562 | 0.15 | **STAT3** | HeLa-S3 | 0.14 |
| IRF3 | HeLa-S3 | 0.22 | **TCF7L2** | HeLa-S3, MCF-7 | 0.19, 0.17 |
| IRF9 | K562 | 0.18 | **TFE3** | K562 | 0.26 |
| ISX | HepG2 | 0.14 | **USF1** | H1, HepG2 | 0.18, 0.15 |
| JUN | HepG2, HeLa-S3 | 0.18, 0.16 | **USF2** | HeLa-S3 | 0.19 |
| JUND | HepG2, HeLa-S3 | 0.17, 0.15 | **ZBTB33** | MCF-7 | 0.14 |
| LEF1 | K562 | 0.13 | **ZKSCAN1** | HeLa-S3, HepG2, K562, MCF-7 | 0.36, 0.25, 0.25, 0.20 |
| MAFF | HepG2, K562, HeLa-S3 | 0.49, 0.41, 0.35 | **ZNF24** | K562, HepG2 | 0.26, 0.15 |
| MAFG | K562, HepG2 | 0.36 , 0.14 | **ZNF274** | HeLa-S3, K562, HepG2 | 5.40,1.35,0.14 |
| MAFK | HepG2, HeLa-S3, MCF-7, H1, K562 | 0.37, 0.25, 0.24, 0.16, 0.15 | **ZNF282** | HepG2 | 0.16 |
| MEF2A | K562 | 0.15 | **ZNF382** | HepG2 | 0.30 |
| MEF2D | K562 | 0.18 | **ZNF460** | HepG2 | 0.13 |
| MITF | K562 | 0.24 | **ZNF652** | HepG2 | 0.14 |

**Supplementary Table 7.** Predicted potential pioneer factors from enrichment analysis using all identified NFRs and NDRs in open chromatin regions. Top 25% transcription factors found by the enrichment analysis by calculating the binding motif enrichment on nucleosomal regions (NRs) compared to nucleosome-depleted regions (NDRs) and are highly expressed (RPKM >=10) in corresponding cell lines.

| TF name | Cell line | Enrichment Score | Expression level (RPKM value) |
| --- | --- | --- | --- |
| ATF2 | HepG2, K562 | 0.65, 0.43 | 35.35, 49.27 |
| ATF4 | K562 | 0.46 | 374.61 |
| ATF7 | HepG2 | 0.59 | 11.26 |
| CEBPB | K562, HepG2, H1, HeLa-S3 | 0.47, 0.26, 0.19,0.25 | 10.53,42.83, 13.88,29.7 |
| CEBPG | K562, HepG2 | 0.52, 0.13 | 33.14, 31.21 |
| ESRRB | K562 | 0.14 | 11.63 |
| FOS | HeLa-S3 | 0.15 | 11.02 |
| FOXA2 | HepG2 | 0.15 | 10.88 |
| FOXA3 | K562 | 0.20 | 12.74 |
| GATA1 | K562 | 0.13 | 125.16 |
| HLF | HepG2 | 0.31 | 180.96 |
| HMBOX1 | K562 | 0.24 | 27.41 |
| IRF3 | HeLa-S3 | 0.22 | 23.49 |
| JUN | HepG2 | 0.18 | 28.16 |
| JUND | HepG2, HeLa-S3 | 0.17, 0.15 | 46.02,14.06 |
| MAFF | HepG2 | 0.49 | 13.95 |
| MAFG | K562, HepG2 | 0.36, 0.14 | 26.12, 21.66 |
| NFATC3 | K562 | 0.14 | 56.33 |
| NFE2 | K562 | 0.18 | 143.18 |
| NFE2L2 | HepG2 | 0.21 | 75.31 |
| NFYB | HeLa-S3 | 0.14 | 21.26 |
| PBX2 | K562 | 0.27 | 49.68 |
| RBPJ | K562 | 0.16 | 22.70 |
| REST | H1, HeLa-S3 | 0.14,0.26 | 17.98,18.56 |
| RFX5 | HeLa-S3 | 0.18 | 57.07 |
| SPI1 | K562 | 0.14 | 12.52 |
| SREBF2 | HeLa-S3 | 0.15 | 23.51 |
| SRF | H1, K562, HepG2 | 0.30, 0.20, 0.14 | 28.78, 28.71, 51.26 |
| STAT3 | HeLa-S3 | 0.14 | 25.54 |
| USF1 | H1, HepG2 | 0.18, 0.15 | 15.91, 16.61 |
| USF2 | HeLa-S3 | 0.19 | 34.93 |
| ZKSCAN1 | HepG2, K562, HeLa-S3 | 0.25, 0.25,0.36 | 49.98, 29.79,14.09 |
| ZNF24 | K562, HepG2 | 0.26, 0.15 | 53.34, 19.30 |
| ZNF274 | K562, HepG2, HeLa-S3 | 1.35, 0.14,5.4 | 27.04, 15.37,13.84 |
| ZNF282 | HepG2 | 0.16 | 11.46 |

**Supplementary Table 8.** Transcription factors that have binding sites enriched on nucleosomal regions located in the differentially open chromatin regions (Enrichment score >=1 and q-value <=0.05).

| TF name | Cell line | Enrichment score | TF name | Cell line | Enrichment Score |
| --- | --- | --- | --- | --- | --- |
| ATF4 | K562 | 1.44 | **MAFF** | HeLa-S3,HepG2,K562 | 1.38,1.25,2.35 |
| BACH1 | K562 | 2.20 | **MAFG** | HepG2,K562 | 1.28,2.79 |
| BCL6 | HepG2 | 1.17 | **MAFK** | K562 | 1.90 |
| CEBPA | HepG2 | 2.95 | **MEF2A** | HepG2,K562 | 1.95,2.51 |
| CEBPB | HeLa-S3,HepG2,K562 | 2.58,1.68,1.26 | **MEF2D** | K562 | 3.11 |
| CEBPD | HepG2 | 2.84 | **MEIS2** | K562 | 1.11 |
| CEBPG | HepG2,K562,MCF-7 | 1.92,1.29,1.35 | **NFATC3** | K562 | 1.76 |
| CUX1 | K562 | 1.29 | **NFE2** | K562 | 2.28 |
| ESRRA | HepG2 | 1.42 | **NFE2L1** | K562 | 2.75 |
| FOS | HeLa-S3,K562,MCF-7 | 1.25,1.91,1.78 | **NFIC** | HepG2 | 1.86 |
| FOSL1 | K562 | 1.89 | **NFIL3** | HepG2 | 1.80 |
| FOSL2 | HepG2,MCF-7 | 1.46,1.76 | **NFIX** | K562 | 1.46 |
| FOXA1 | HepG2,K562,MCF-7 | 2.76,4.27,2.78 | **NR2F1** | HepG2,K562 | 1.61,1.08 |
| FOXA2 | HepG2 | 2.26 | **NR2F2** | HepG2,K562 | 2.70,1.75 |
| FOXA3 | HepG2,K562 | 2.01,2.61 | **NR4A1** | K562 | 1.87 |
| FOXK2 | K562 | 1.35 | **NR5A1** | HepG2 | 1.81 |
| FOXP1 | HepG2 | 2.38 | **ONECUT1** | HepG2 | 1.72 |
| GATA1 | K562 | 6.15 | **ONECUT2** | HepG2 | 2.33 |
| GATA2 | HepG2,K562 | 4.51,6.39 | **PBX2** | HepG2,K562 | 1.70,1.87 |
| GATA3 | MCF-7 | 2.14 | **PPARG** | HepG2 | 1.13 |
| GATA4 | HepG2 | 2.54 | **PRDM1** | HeLa-S3 | 1.60 |
| GFI1 | HepG2 | 1.14 | **RARA** | HepG2 | 1.95 |
| HLF | HepG2 | 1.41 | **RUNX1** | K562 | 2.93 |
| HMBOX1 | K562 | 1.55 | **RXRB** | HepG2 | 2.03 |
| HNF1A | HepG2 | 4.09 | **SOX13** | HepG2 | 2.43 |
| HNF1B | HepG2 | 1.55 | **SRF** | K562 | 2.05 |
| HNF4A | HepG2 | 3.64 | **STAT3** | HeLa-S3 | 4.25 |
| HNF4G | HepG2 | 3.32 | **TCF7** | HepG2,K562 | 3.51,1.30 |
| IKZF1 | K562 | 1.35 | **TCF7L2** | HepG2,MCF-7 | 2.73,1.42 |
| IRF1 | K562 | 1.31 | **TEAD1** | HepG2 | 1.36 |
| IRF9 | K562 | 2.57 | **TEF** | HepG2 | 1.61 |
| JUN | HeLa-S3 | 1.22 | **THRB** | HepG2 | 2.15 |
| JUNB | K562 | 2.06 | **ZNF24** | K562 | 1.13 |
| JUND | HeLa-S3,HepG2,K562 | 2.27,1.44,1.97 | **ZNF652** | HepG2 | 1.30 |
| LEF1 | K562 | 4.48 |  |  |  |

**Supplementary Table 9.** Predicted potential pioneer factors from enrichment analysis using the NRs located in differentially open chromatin regions and NDRs located in conserved open chromatin regions. Transcription factors that have binding sites enriched on nucleosomal regions in the differentially open chromatin regions (Enrichment score >=1 and q-value <=0.05) and are highly expressed (RPKM >=10) in corresponding cell lines.

| TF name | Cell line | Enrichment score | Expression level (RPKM value) |
| --- | --- | --- | --- |
| ATF4 | K562 | 1.44 | 374.61 |
| BACH1 | K562 | 2.20 | 21.38 |
| BCL6 | HepG2 | 1.17 | 28.63 |
| CEBPA | HepG2 | 2.95 | 33.37 |
| CEBPB | HeLa-S3,HepG2,K562 | 2.58,1,68,1.26 | 29.70,42.83,10.53 |
| CEBPD | HepG2 | 2.84 | 26.11 |
| CEBPG | HepG2,K562 | 1.92,1.29 | 31.21,33.14 |
| ESRRA | HepG2 | 1.42 | 16.10 |
| FOS | HeLa-S3 | 1.25 | 11.02 |
| FOSL2 | HepG2 | 1.46 | 19.26 |
| FOXA1 | HepG2 | 2.76 | 32.38 |
| FOXA2 | HepG2 | 2.26 | 10.88 |
| FOXA3 | HepG2,K562 | 2.01,2.61 | 57.20, 12.74 |
| FOXK2 | K562 | 1.35 | 22.64 |
| GATA1 | K562 | 6.15 | 125.16 |
| GATA2 | K562 | 6.39 | 18.36 |
| GATA4 | HepG2 | 2.54 | 13.42 |
| HLF | HepG2 | 1.41 | 180.96 |
| HMBOX1 | K562 | 1.55 | 27.41 |
| HNF4A | HepG2 | 3.64 | 72.69 |
| IKZF1 | K562 | 1.35 | 14.53 |
| JUNB | K562 | 2.06 | 19.14 |
| JUND | HeLa-S3,HepG2,K562 | 2.27,1.44,1.97 | 14.06,46.02,41.06 |
| LEF1 | K562 | 4.48 | 13.49 |
| MAFF | HepG2 | 1.25 | 13.95 |
| MAFG | HepG2,K562 | 1.28,2.79 | 21.66,26.12 |
| MEF2A | HepG2 | 1.95 | 14.56 |
| NFATC3 | K562 | 1.76 | 56.33 |
| NFE2 | K562 | 2.28 | 143.18 |
| NFE2L1 | K562 | 2.75 | 28.47 |
| NFIL3 | HepG2 | 1.80 | 51.02 |
| NR2F1 | HepG2 | 1.61 | 28.15 |
| NR5A1 | HepG2 | 1.81 | 28.13 |
| PBX2 | HepG2,K562 | 1.70,1.87 | 37.77,49.68 |
| PPARG | HepG2 | 1.13 | 26.05 |
| RXRB | HepG2 | 2.03 | 34.15 |
| SRF | K562 | 2.05 | 28.71 |
| STAT3 | HeLa-S3 | 4.25 | 25.54 |
| TEAD1 | HepG2 | 1.36 | 13.31 |
| ZNF24 | K562 | 1.13 | 53.34 |

**Supplementary Table 10.** Three groups of known pioneer factors for validation of enrichment analysis.

| Test set 1: 32 known pioneer factors | FOXA1, FOXA2, FOXA3, GATA1, GATA2, GATA3, GATA4, CEPBA, CEBPB, NFYA, NFYB, NFYC, ESRRB, POU5F1, KLF4, NEUROD1, TP53, FOS, FOSL1, FOSL2, MAFG, MAFF, MAFK, JUN, JUNB, JUND, ATF2, ATF3, ATF4, ATF6, ATF7, SPI1 |
| --- | --- |
| Test set 2: 11 known pioneer factors with essential roles in cell differentiation | FOXA1, FOXA2, FOXA3, GATA1, GATA2, GATA3, GATA4, CEPBA, CEBPB, NEUROD1, SPI1 |
| Test set 3: 7 known pioneer factors for the maintenance of embryonic stem cell or reprogramming of somatic cells into induced pluripotent stem cells | NFYA, NFYB, NFYC, ESRRB, NEUROD1, KLF4, POU5F1 |

**Supplementary Table 11.** Performance of enrichment scores in the classification of pioneer factors.

| NFR and NDR | Validation Set | Highly Expressed TFs | ROC-AUC | | pr-ROC-AUC | MCC (maximum value) | Percentage of TFs as True Positives |
| --- | --- | --- | --- | --- | --- | --- | --- |
| All | Test Set 1 | No | 0.69 | 0.33 | | 0.31 | 0.17 |
|  | Test Set 1 | Yes | **0.71** | **0.37** | | **0.31** | 0.2 |
|  | Test Set 2 | No | 0.77 | 0.1 | | 0.22 | 0.04 |
|  | Test Set 2 | Yes | 0.76 | 0.13 | | 0.22 | 0.06 |
| Differentially and conserved open chromatin regions | Test Set 1 | No | 0.71 | 0.34 | | 0.28 | 0.16 |
|  | Test Set 1 | Yes | 0.71 | 0.41 | | 0.33 | 0.2 |
|  | Test Set 2 | No | 0.89 | 0.41 | | 0.42 | 0.04 |
|  | Test Set 2 | Yes | **0.92** | **0.45** | | **0.49** | 0.06 |

**Supplementary Table 12.** List of TFs that have been previously suggested *as potential pioneer factors* and/or *nucleosome binders* from the literature.

| TF name | Function | Reference |
| --- | --- | --- |
| LEF1 | able to bind with HIV-1 nucleosome core particles | ^1^ |
| HNF4A | a potential pioneer factor in remodeling the active chromatin landscape in the liver | ^2^ |
| PBX2 | one PBC transcription factor which could have pioneer activity | ^3^ |
| NR5A1 | considered as a pioneer transcription factor for Leydig cell development and function | ^4^ |
| NFE2 | interact with the cognate motif on the nucleosome before chromatin is remodeled | ^5^ |
| CUX1 | specifically interacts with its recognition motif in a nucleosomal context without displacing the nucleosome core | ^6^ |
| FOXK2 | possibly acts as an early pioneer factor in gene activation during cell differentiation | ^7^ |
| NRF1 | predicted on theoretical bases to have pioneer action and trigger chromatin access (DNase sensitivity) only if its DNA-binding site is unmethylated | ^8^ |
| RFX5 | displace nucleosomes | ^9^ |
| CREB1 | bind to inaccessible chromatin regions | ^9^ |
| FOXO4 | Potential pioneer factors in cancer development | ^8^ |
| CLOCK | CLOCK:BMAL1 DNA binding promotes rhythmic chromatin opening | ^10^ |

**Supplementary Table 13.** Predicted potential pioneer factors from the association analysis between the binding motif profiles and nucleosome occupancy values. A list of TFs with statistically significant positive correlations between the TF binding motif profiles and nucleosome occupancy levels (p-value <=0.05). TF expression level data is not available for MCF-7 cell line.

| TF name | Cell Line | PCC | Expression level |
| --- | --- | --- | --- |
| ZKSCAN1 | HeLaS3, K562, MCF7, HepG2 | 0.75,0.64,0.63, 0.34 | 14.1,29.8, NA, 50.0 |
| ESR1 | MCF7 | 0.58 | NA |
| NFATC3 | K562 | 0.56 | 56.3 |
| ZBTB7B | MCF7, HepG2 | 0.55,0.37 | NA, 15.0 |
| MAX | H1 | 0.51 | 12.4 |
| TBX2 | HepG2 | 0.50 | 2.2 |
| NFIB | MCF7 | 0.48 | NA |
| USF1 | H1 | 0.46 | 15.9 |
| CUX1 | MCF7, K562 | 0.41,0.08 | NA, 2.4 |
| ZNF274 | HepG2 | 0.37 | 15.4 |
| SREBF1 | MCF7 | 0.36 | NA |
| NFYB | HeLaS3 | 0.33 | 21.3 |
| ZNF24 | MCF7, K562 | 0.32,0.08 | NA, 53.3 |
| NR2F2 | MCF7 | 0.31 | NA |
| ZNF282 | K562 | 0.30 | 17.7 |
| RELA | K562 | 0.30 | 8.9 |
| USF2 | HeLaS3 | 0.28 | 34.9 |
| ESRRA | MCF7 | 0.28 | NA |
| POU5F1 | H1 | 0.26 | 183.0 |
| NFKB2 | HepG2 | 0.22 | 11.1 |
| BACH1 | H1 | 0.22 | 6.3 |
| SREBF2 | HeLaS3 | 0.21 | 23.5 |
| TEAD4 | H1 | 0.20 | 44.0 |
| MIXL1 | HepG2 | 0.18 | 9.76 |
| NR4A1 | K562 | 0.15 | 0.8 |
| E2F6 | H1 | 0.15 | 3.1 |
| RARA | HepG2 | 0.14 | 2.7 |
| CEBPB | HeLaS3 | 0.13 | 29.7 |
| SMAD5 | K562 | 0.12 | NA |
| NFYA | HeLaS3 | 0.11 | 12.5 |
| HMBOX1 | K562 | 0.11 | 27.4 |
| ZNF460 | HepG2 | 0.11 | 12.7 |
| CTCF | H1 | 0.10 | 30.7 |
| EGR1 | H1 | 0.10 | 4.80 |
| GATA3 | MCF7 | 0.09 | NA |
| ATF2 | K562 | 0.08 | 49.3 |
| TCF7L2 | MCF7 | 0.07 | NA |

**Supplementary Figure 1.** Identification of differentially open and conserved open chromatin regions between H1 embryonic cell line and any other differentiated cell line. Open chromatin regions are shown as lines.

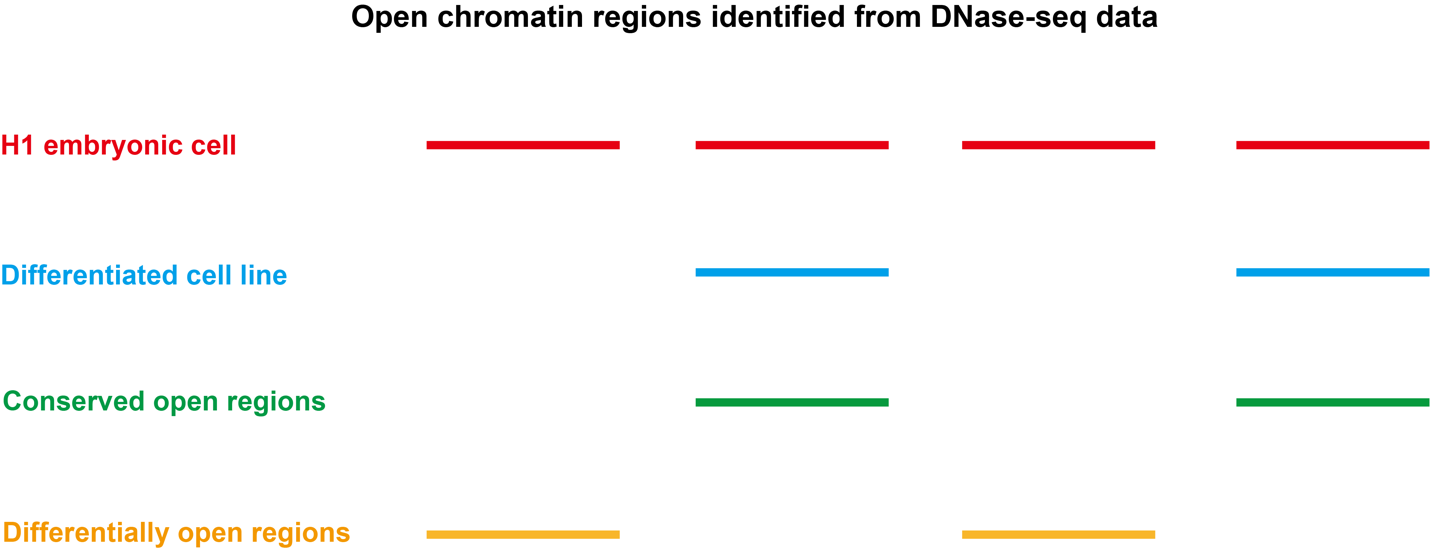

**Supplementary Figure 2.** Identification of dinucleotide patterns of nucleosomal DNA (MCF7 cell line is shown as a representative case).

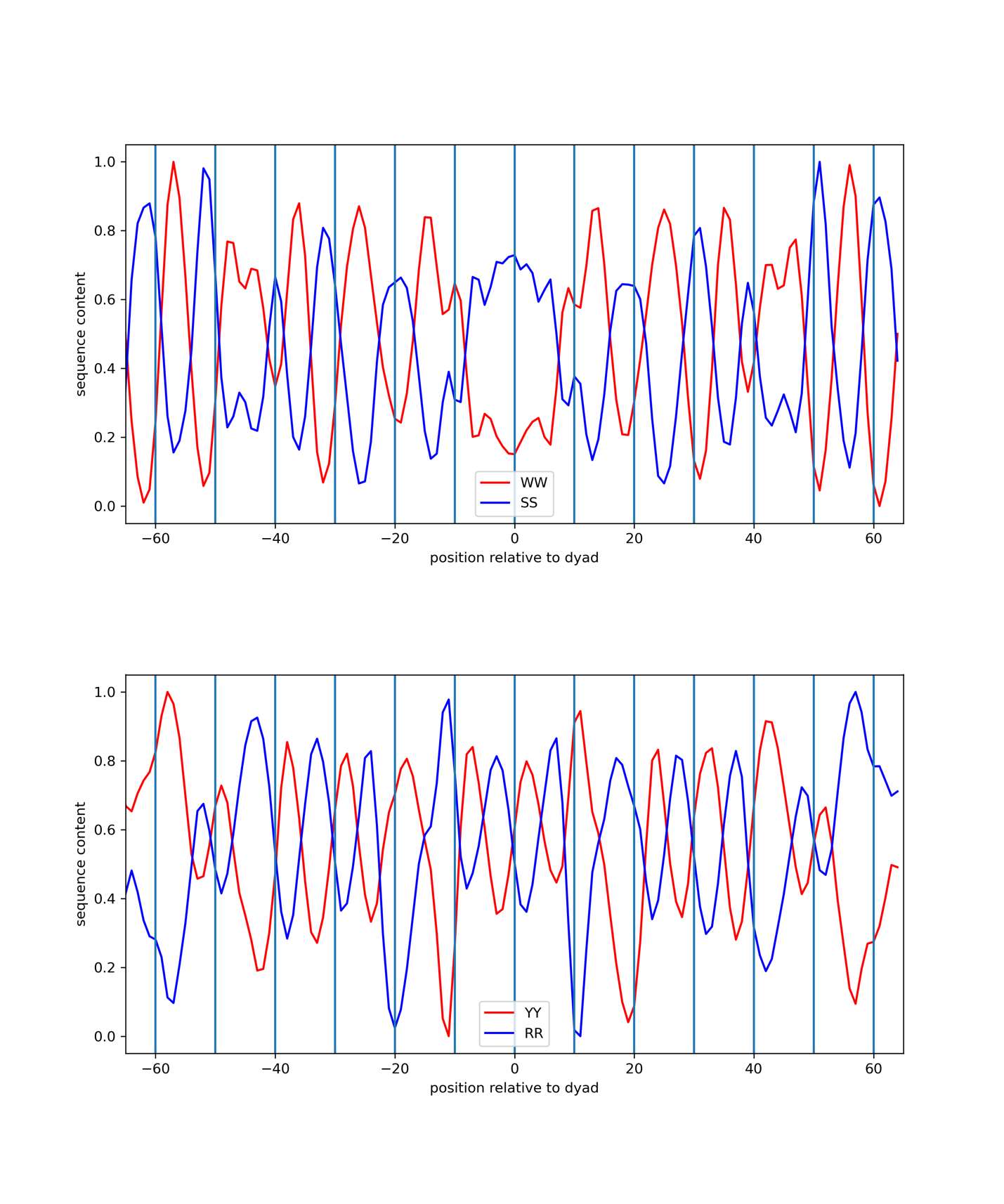

**Supplementary Figure 3.** Motif enrichment analysis of different TFs using NDRs located in open chromatin regions and all identified NRs. 32 Known PTFs from Supplementary Table 5 and non-PTFs are shown with red and light blue colors. Mann–Whitney U test is performed under the null hypothesis that the mean of enrichment scores of PTFs is larger than for other TFs. a) and b) show the ranking of all TFs and significantly expressed TFs by their enrichment score. ** - p-value < 0.001.

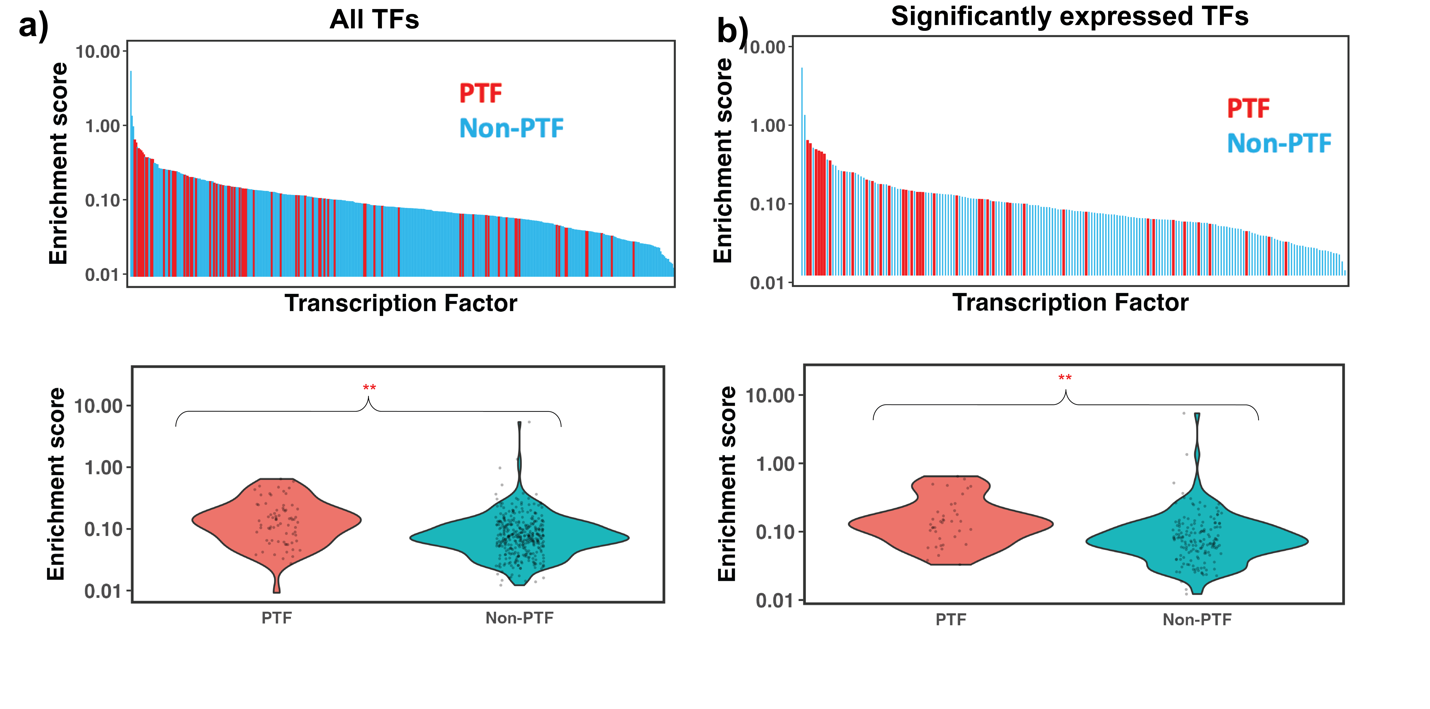

**Supplementary Figure 4.** Comparison of the enrichment scores of pioneer factors for the maintenance of embryonic stem cell or reprogramming of somatic cells into induced pluripotent stem cells (Test set 3, highlighted as red) with other TFs. Pioneer factors in Test set 3 were strongly depleted at nucleosomes and showed significantly lower enrichment scores compared with other TFs.

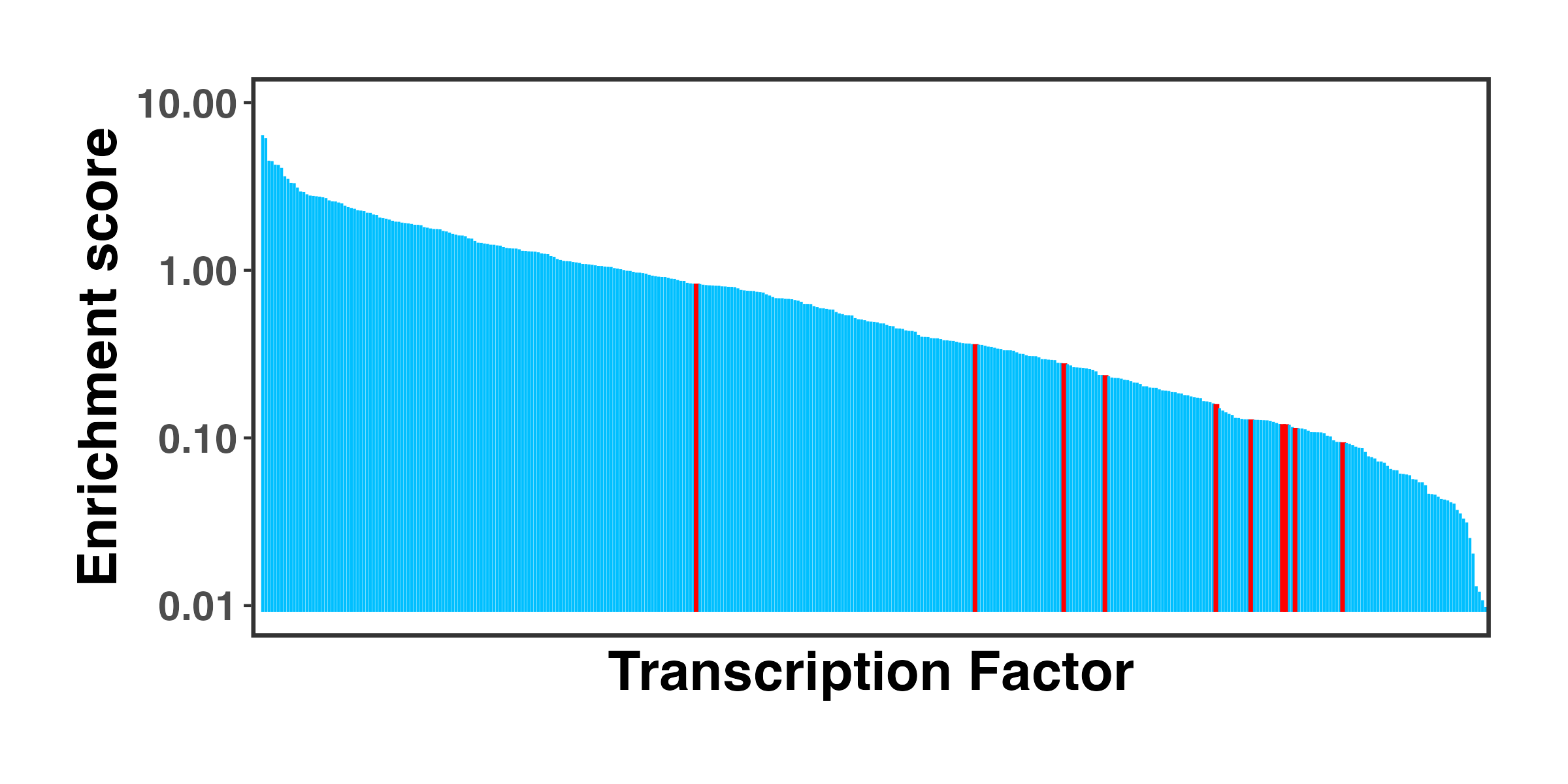

**Supplementary Figure 5.** Enrichment analysis of TFs by redefining differentially open regions as those closed in differentiated cell lines and open in H1 embryonic cell line. Known pioneer factors with essential roles in cell differentiation (Test set 2) and pioneer factors critical for the maintenance of embryonic stem cell or reprogramming of somatic cells into induced pluripotent stem cells (Test set 3) are indicated by squares and triangles, while other TFs are shown as circles. Colors corresponds to false discovery rate (FDR) q-values. ESSRB and Yamanaka pioneer transcription factor POU5F1 (OCT4) from Test set 3 showed significantly higher enrichment scores.

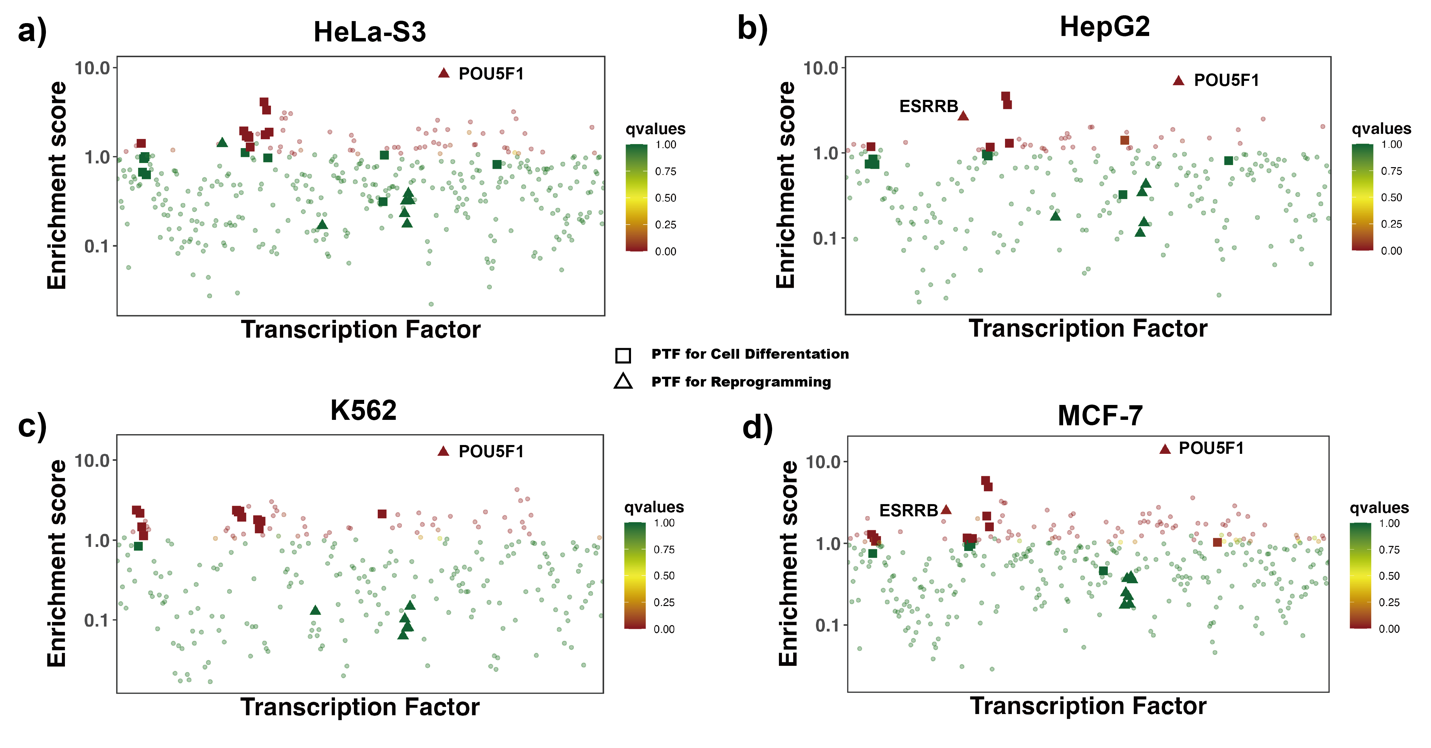

**Supplementary Figure 6.** TF motif enrichment score is used to distinguish 11 known PTFs with essential roles in cell differentiation (Test set 2) from other TFs. Receiver operating curve (ROC) and precision-recall (PR) curve analysis are performed. Enrichment scores are calculated using the NRs and NDRs from a) differentially and conserved open chromatin regions and b) differentially and conserved active enhancer regions.

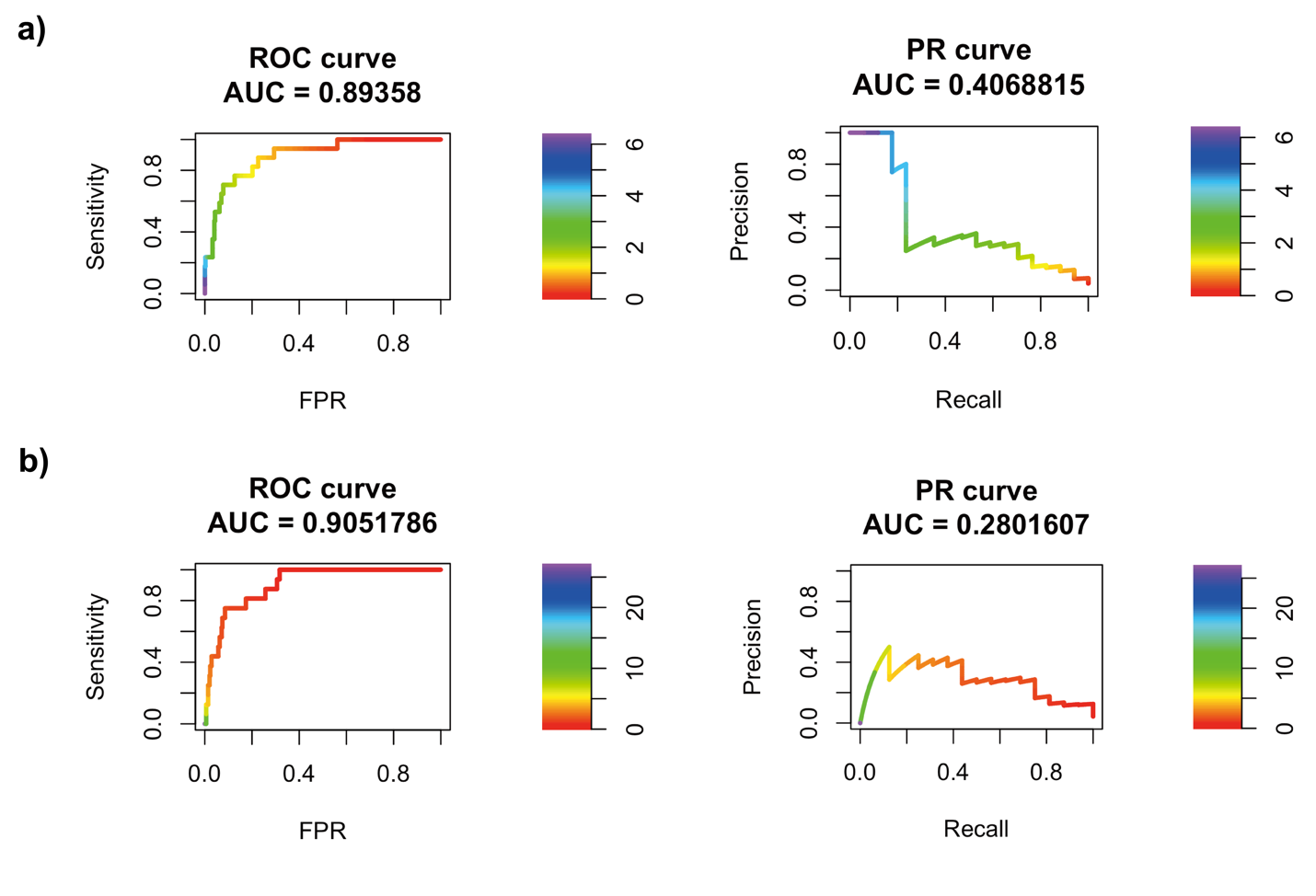

**Supplementary Figure 7.** TF motif enrichment score is used to distinguish 11 known PTFs with essential roles in cell differentiation (Test set 2) from other TFs. Receiver operating curve (ROC) analyses of motif enrichment scores are performed using different thresholds in defining the differentially and conserved open chromatin regions.

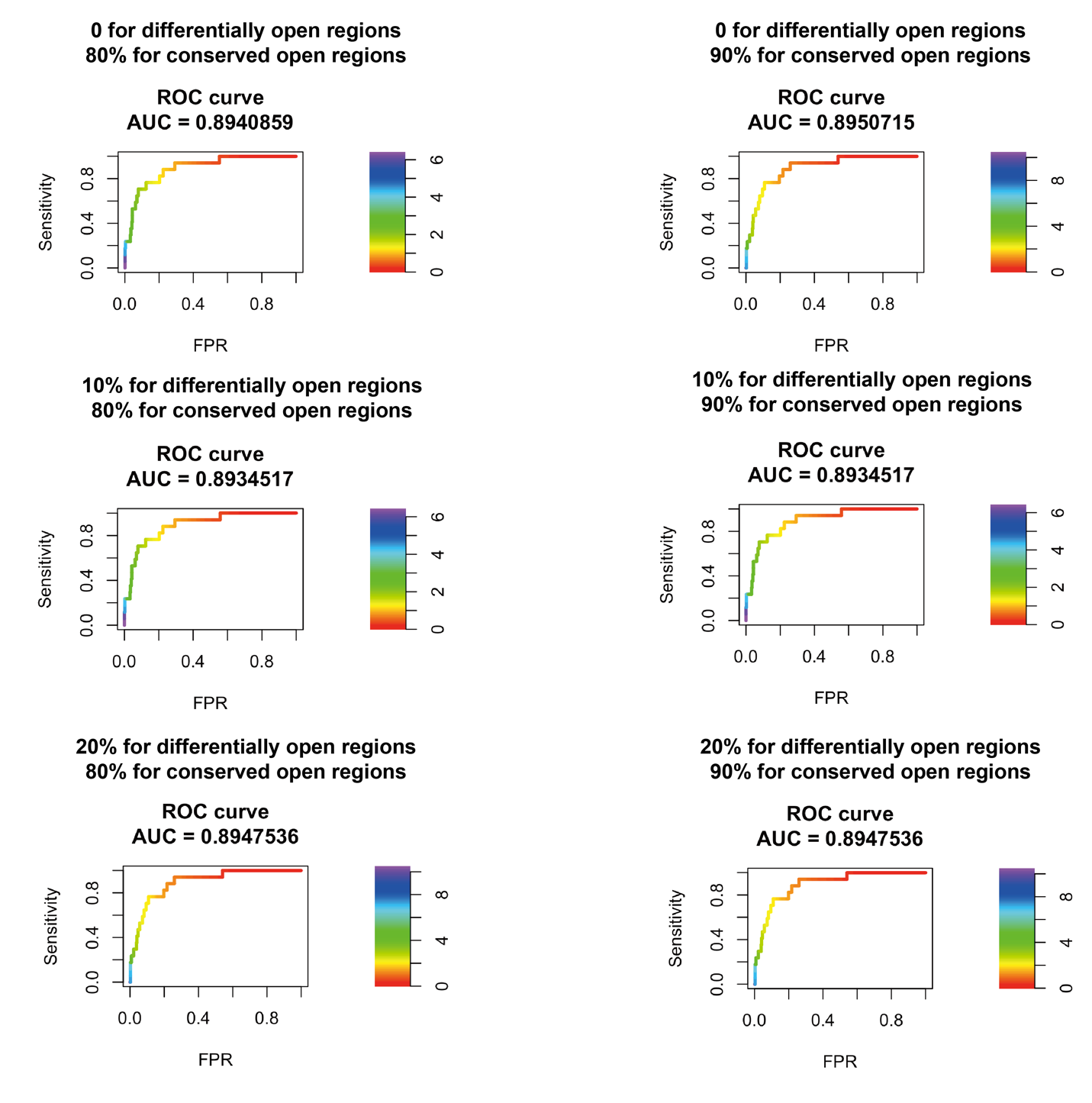

**Supplementary Figure 8.** A liner regression model describing the relationship between calculated TF end/dyad binding ratio ($\tau_{end/dyad}$) from our study and the EMI intensity and EMI penetration values from recent nucleosome NCAP–SELEX experiments^11^. A Pearson correlation coefficient is shown.

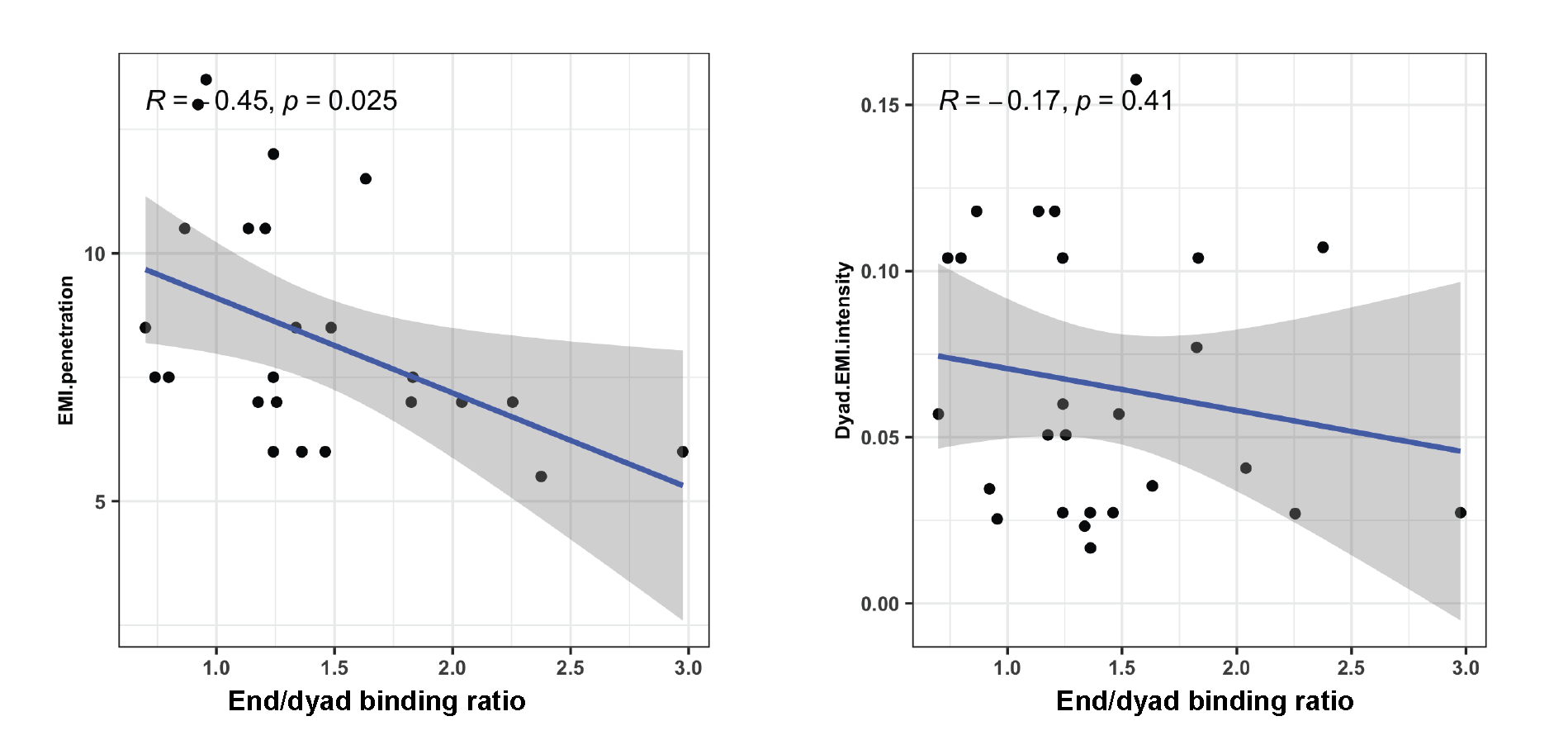

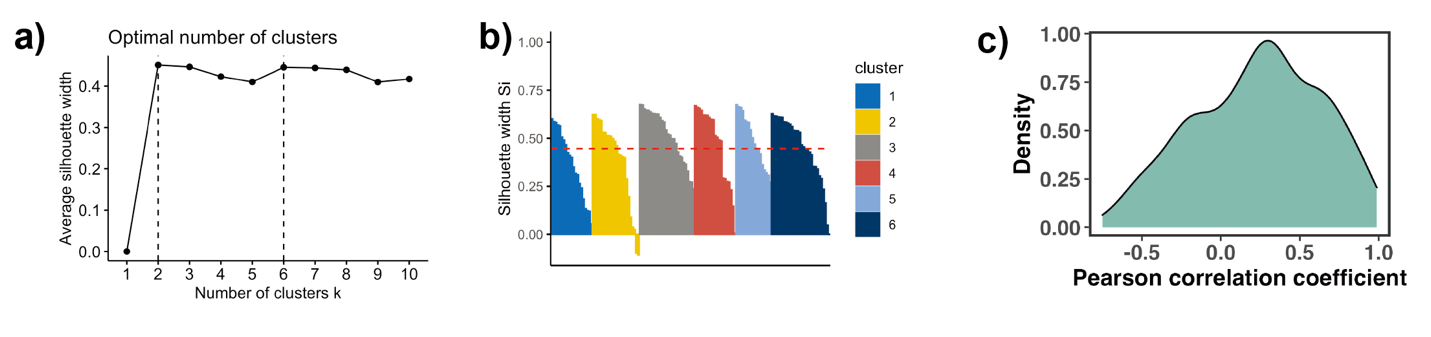
**Supplementary Figure 9.** **a)** Variation of the average silhouette width with the number of clusters in k-medoids clustering of TF binding profiles. Six is selected as an optimal number of clusters. **b)** Distribution of silhouette width for all points (S_i_) in each cluster. Each point with S_i_ <=0.25 is considered as outlier and excluded from the analysis. **c)** Pearson correlation coefficients (PCCs) of binding motif profiles between two symmetrical nucleosomal halves (define SHL) of all TFs. TFs with PCC values less than 0.4 and the total number of base pairs of binding motifs in NRs less than 500 are excluded from the analysis.
